# Scaling up: understanding movement from individual differences to population-level dispersal

**DOI:** 10.1101/2021.01.04.425125

**Authors:** Allan H. Edelsparre, Trevor J. Hefley, Marco A. Rodríguez, Mark J. Fitzpatrick, Marla B. Sokolowski

**Affiliations:** Department of Ecology & Evolution, University of Toronto, Toronto ON, M5S 3B2, Canada; Department of Cell & Systems Biology, University of Toronto ON, M5S 3G5, Canada; Integrative Behaviour and Neuroscience Group, Department of Biological Sciences, University of Toronto Scarborough, Toronto ON, M1C 1A4, Canada; Department of Statistics, Kansas State University, Manhattan KS, 66506, USA; Départment des sciences de l’environnement, Université du Québec à Trois-Rivières,C.P. 500, Trois-Rivières, Québec G9A 5H7, Canada; Child and Brain Development Program, Canadian Institute for Advanced Research (CIFAR), Toronto, Ontario, M5G 1M1, Canada

**Keywords:** Movement ecology, dispersal, behaviour, individual variation, ecological diffusion, advection, *foraging* gene, *Drosophila melanogaster*.

## Abstract

Dispersal is fundamental to life on our planet. Dispersal facilitates colonization of continents and islands. Dispersal mediates gene flow among populations, and influences the rate of spread of invasive species. Theory suggests that individuals consistently differ in dispersal propensity, however determining the relative contributions of environmental factors to individual and population-level dispersal, represent a major challenge to understand the spread of organisms. To address this, we conducted a field experiment using *Drosophila melanogaster.* As proxies for individuals with different dispersal propensities, we used wildtype strains of flies with natural variants of the *foraging* gene, known to influence dispersal in laboratory and field experiments. These included flies with *for*^s^ alleles known to be less dispersive, flies with the *for*^R^ alleles which are more dispersive flies as well as an outbred population established from field collected flies. We released approximately 6000 flies of each strain in an experimental arena (100 m × 100 m) in the field and our recaptures were used to determine dispersal of flies over time. To estimate environmental effects on dispersal, we measured temperature, wind direction and wind speed. Using partial-differential equations we combined ecological diffusion with advection to estimate dispersal rates and responses to wind. We found that temperature effects elicited a similar response in high and low dispersal lab strains with dispersal rate increasing with temperature most rapidly at temperatures above 18°C. This was in contrast to outbred flies which remained unresponsive to temperature changes. We also detected a response to wind with advection rates increasing linearly with wind speed for all flies in general. Our results suggest that response to temperature and wind can minimize known differences in behavioural predispositions to disperse. Our results also suggest that the direction and magnitude of wind may play a key role in the colonization and distribution of fly populations. Our findings therefore have implications for forecasting the spread of pests and invasive species as well as pathogens and vectors of disease. Our findings further contribute to the understanding of how the environment can modify behavioural predispositions and to influence population-level dispersal in fly populations in particular and insect species in general.

## 1. Introduction

Dispersal plays a fundamental role in the evolutionary and ecological processes that govern life on our planet. From an evolutionary perspective, dispersal influences range expansion and rates of diversity (Bocxlaer et al. 2010) as well as adaptive radiations from mainland continents to islands. An example of the latter is the dispersal of anole lizards across the Greater Antilles that has facilitated the repeated evolution of similar ecomorphs on separate islands (Losos et al. 1998). Additionally, the evolution of Darwin’s finches and Hawaiian honeycreepers involved a single dispersal event from the mainland that facilitated the subsequent evolution of many different forms across the archipelagos (Grant 1981; Freed et al. 1987). From an ecological perspective, dispersal influences rates of birth, death and immigration among populations and communities (Hanski and Gilpin 1991). Dispersal mediates gene flow among populations, and is an important consideration in studies of urbanized and fragmented landscapes (Cote et al. 2017, Edelsparre et al. 2018). Finally, dispersal can also influence the rate and severity of the spread of pathogens as well as invasive species (Kot et al. 1996).

Individuals within a population can disproportionally influence dispersal. Invasive cane toads from older established areas move slower and have reduced reproductive rates relative to individuals at the leading front of the invasion (Phillips 2009) which have longer legs, move faster and in straighter paths (Phillips et al. 2006). Increased dispersal rates of individuals which are at the leading front of invasions have also been observed in butterflies (Hill et al. 1999), aphids (Lombeart et al. 2014), and fish (Myles-Gonzalez et al. 2015). Understanding how individual variation contributes to population dispersal can have important implications for our ability to model and predict population level dispersal. A lack of understanding of the factors that affect dispersal can result in an underestimation of the spread of invasive species and the capacity for species to reclaim or colonize new habitats (Kot et al. 1996; Saastamoinen et al. 2018). This, in turn, limits our ability to make accurate ecological forecasts (Clarke et al. 2001), which affects our ability to manage the effect of spread of invasive species on ecosystems that are being invaded or colonized (Hastings et al. 2005; Gomez-Uchida et al. 2018).

Elucidating the underlying factors that contribute to dispersal is a major challenge for the field of movement ecology (Nathan et al. 2008; Cote et al. 2017). Several morphological, physiological, and behavioural traits are linked with variation in dispersal. For example, insect wing dimorphisms that affect dispersal ability are common (e.g. pea aphid: Zera and Deno 1997; rice plant hoppers: Brisson 2010). Behavioural traits have also been linked with dispersal in a wide range of animal taxa, including birds, fishes, and lizards (Réale et al. 2007; Cote et al. 2017). Individuals have been shown to differ in their dispersal propensity depending on whether they are aggressive (Duckworth and Badyaev 2007), are risk takers (Edelsparre et al. 2013), or are sociable (Cote and Clobert 2007). In rare cases, genes that underlie the link between these behaviours and dispersal have been identified (Korsten et al. 2013; Edelsparre et al. 2014) offering unique insights into the potential mechanisms contributing to individual differences in dispersal. However, how such factors play out in nature to influence dispersal at the population-level is poorly understood (Gurarie et al. 2009).

There are three main hypotheses through which population-level variation in dispersal might arise. First, individuals within populations can differ in their predispositions to disperse if genetic differences between individuals affect dispersal directly or via differences in morphology, physiology and/or behaviour, such as those mentioned above (Sastaamoinen et al. 2018). Under this hypothesis we would expect populations to display consistent individual differences in dispersal across multiple environmental contexts (changes in climate and/or landscape). Such consistent differences could exist even if groups or individuals with different dispersal predispositions responded similarly (parallel responses) and/or differently (divergent responses) to environmental change. Second, variation in dispersal could arise largely in response to environmental factors. This could be if dispersal predispositions are absent or if the environmental effects are strong enough to minimize dispersal predispositions. Under this scenario we would expect to see a general population response to the environment and with changes in the environment driving changes in dispersal (each disperser is a random draw from the population). Finally, variation in dispersal could arise through a combination of the above possibilities, including conditions ranging between individual (e.g. dispersal predispositions) and environmentally driven dispersal. Detecting complex relationships between key factors such as those outlined above would require the implementation of dynamic models that are capable of estimating the relative contribution of each factor on the movement process (Hefley et al. 2017; Hooten and Hefley 2019).

In the following study we set out to address these three hypotheses by combining individual-level predictors and environmental data in dynamic models designed to assess the relationship between each factor over time. In our experiment, we used the common fruit fly, *Drosophila melanogaster,* a convenient organism to model dispersal in general and to investigate individual-level predictors of dispersal in particular. To accomplish this we first produced a large outbred study population from field collected flies. Secondly, we incorporated individual-level predictors of dispersal by using *Drosophila melanogaster* that carry variants of a gene known to influence differences in the propensity of the adult fly to disperse (Edelsparre et al. 2014; Edelsparre et al. 2018). This particular *Drosophila* system consists of two strains of flies that differ in several movement related behaviours both as larvae and adults and these differences are mediated by natural variation in the *foraging* (*for*) gene (Osborne et al. 1987; de Belle et al. 1993; Pereira and Sokolowski 1993; Edelsparre et al. 2014). Individuals that carry *for*^R^ alleles (rovers) are active foragers as larvae and more dispersive as adults while individuals that carry *for*^s^ alleles (sitters) tend to be less active foragers as larvae and less dispersive as adults. Recent laboratory experiments demonstrated environmentally dependent plasticity in dispersal propensity of the rover and sitter variants of the *foraging* gene (Anreiter and Sokolowski 2019, Edelsparre et al. 2020). Consequently, the rover and sitter strains were used to examine how different dispersal propensities may interact with the environment to produce population-level dispersal and the outbred population was used to examine how a population with multiple dispersers respond to the environment.

Temperature and wind are two important environmental factors that influence insect activity and movement in nature (Glick 1942; Taylor 1963; McManus 1988). Work on several insect species suggests that there are critical temperatures below which insects will not initiate dispersal (Taylor 1963) and even lower temperatures beyond which insects are not able to sustain flight (Cockbain 1961). In general, insect dispersal will increase with temperature, likely with some optimal temperature range conducive to movement. However, whether such a response can be linked with dispersal predispositions remain largely unexplored. Although, wind direction and speed are thought to be critical to insect flight the behavioural response to wind is likely not a simple one (McManus 1988). Since the early 1920’s, when large numbers of insects were first recorded in the atmospheric convective layer (Glick 1939), researchers held the view that aerial dispersal was entirely dependent on weather conditions (e.g. passive dispersal; McManus 1988). Most empirical data since then suggested that there is a behavioural component to dispersal. For example, using radars to quantify airflow and movement of small insects in the atmosphere, Wainwright et al. (2017) detected movement velocities independent of airflow. Work on flies is consistent with the findings of Wainwright et al. (2017). Desert species of *Drosophila* (*D. mimica*, *D. nigrospiracula* and *D. mojavensis*) disperse both up and down wind and this behaviour is dependent on food availability (Richardson and Johnston 1975; Markow and Castrezana 2000). In a recapture experiment, Coyne et al. (1982) released groups of *D. melanogaster* and its sister species *D. simulans* in the Death Valley and were able to recapture a proportion of them at an oasis several miles from the release site even though the flies faced cross winds during their flight. Although none of the studies on *Drosophila* explicitly tested wind effects on dispersal the results suggest that, for these flies, dispersal is not entirely passive. As is the case for the effects of temperature on dispersal, it is unknown whether individual flies respond similarly to wind effects or whether there is a behavioural predisposition to respond differently to wind speed and direction.

Here we examined the effects of temperature, and wind direction and speed on the dispersal behaviour of rover and sitter *D. melanogaster* as well as on an outbred population of flies released in the field using a mark/release/recapture experiment. Because flies do not exhibit dispersal behaviour in a strict sense (e.g. departure, settlement), but rather move while foraging, searching for mates, avoiding predators etc. we defined any movement away from a release site as dispersal. This idea fits well with dispersal defined in its simplest form as any movement by individuals leading to spatial spread with potential for genetic mixing (Ronce 2007). We quantify dispersal as the rate at which this spread occurs (see analysis below). To investigate how the dispersal behaviour of rover, sitter and the outbred strain of flies respond to temperature and wind we monitored the weather during the experiment. We explicitly evaluated the general prediction that dispersal increases with temperature and whether our more (rover) and less (sitter) dispersive strains differed from our outbred population. We also explicitly tested whether flies use wind to disperse and we used a novel approach to detect the tendency for flies to disperse either up or down wind in the field. We also determined whether the strains differed in their dispersal in response to wind, both in terms of direction and velocity. Combining individual predictors of dispersal with temperature and wind allowed us to evaluate not only the interaction between individual dispersal propensity and the environmental factors, but also the relative contribution of each factor in population-level dispersal in nature.

## 2. Materials and methods

### 2.1 Fly lines

To evaluate how strain-differences in behaviour influence dispersal of flies in the field we used two inbred strains (rover- *for*^R^ and sitter- *for*^s^) and one outbred strain of flies. The effect of *for* on the dispersal strategy of the rover and sitter inbred strains was documented in Edelsparre et al. (2014) and Edelsparre et al. (2018). The outbred population was established from 92 iso-female lines originally collected in Sudbury, Ontario, Canada (501484.97 E, 5143198.65 N UTM) on August 12, 2012 by Thomas Merritt. Several months prior to the commencement of the field experiment each of the 92 lines were randomly assigned to one of 6 170mL sponge-topped plastic *Drosophila* bottles (4 bottles each containing 15 different lines and 2 bottles each containing 16 lines) containing 40 mL of media as described below and maintained as stocks. Two months prior to the field experiment the six bottles were transferred to population cages and flies were allowed to mix for one generation in 16 open bottles inside the population cage. Hereafter the 16 bottles were extracted from the cage, and brooded over for two more generations before a fresh generation of flies was used in the field experiment.

### 2.2 Experimental field

To quantify the movement of flies in the field, we prepared a 100 m by 100 m large experimental arena in an open meadow at the ***rare*** **Charitable Research Reserve** west of Cambridge, Ontario, Canada (553195.00 E, 4802841.00 N UTM) that was recently converted from agricultural land to a nature reserve (Fig. 1a). The field is flat with a sloping gradient of zero from corner to corner and mainly consists of vegetation such as goldenrods (*Solidago* spp.), asters (*Symphyotrichum* spp.), thistles (*Circium* spp), and common milkweed (*Asclepias syriaca*). Vegetation reached an average height of approximately 40-50 cm and the density was largely uniform across the entire field (personal observation).

**Figure 1:**
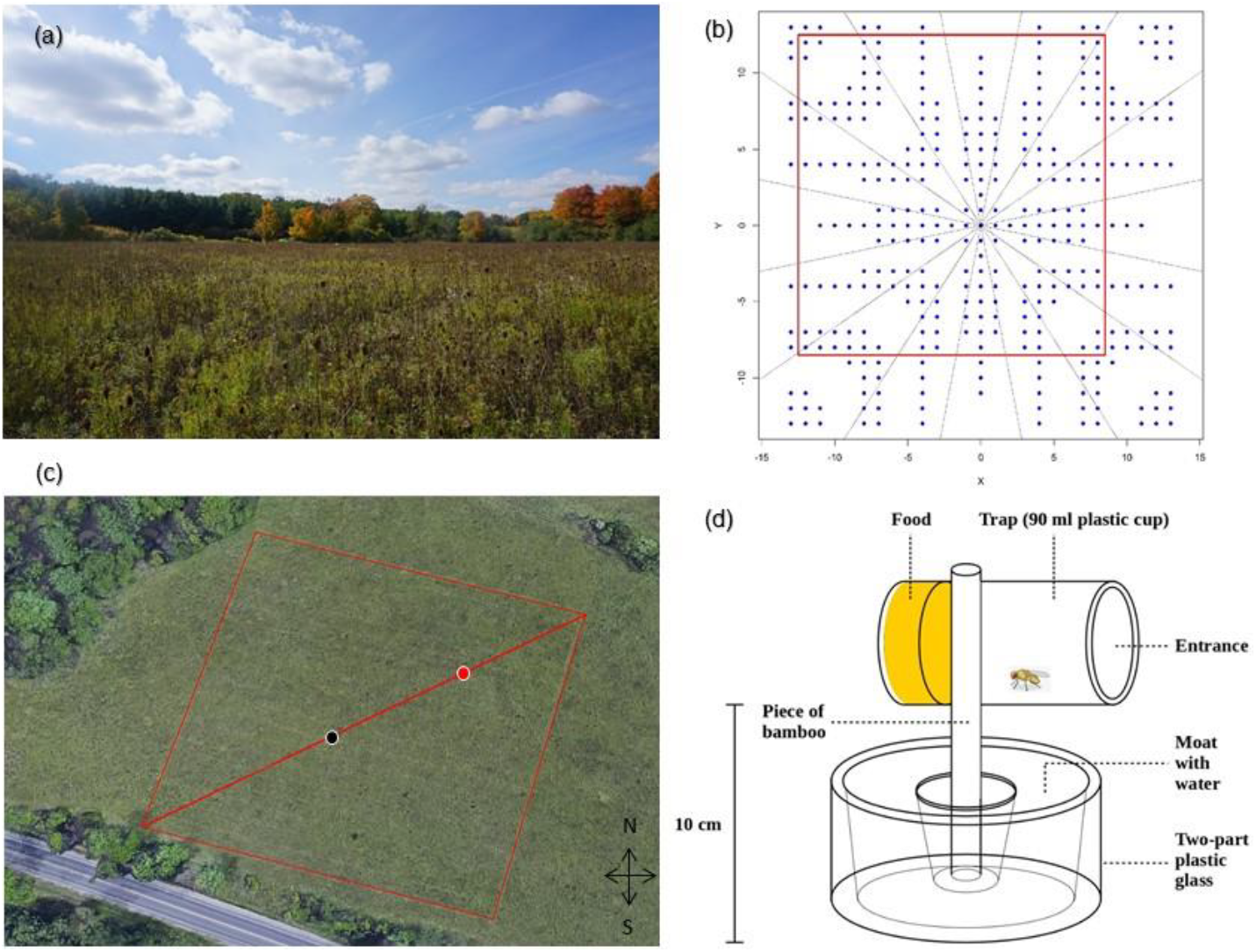
(a) The experimental field looking towards the South-West towards Blair Road, Cambridge, ON, Canada. The site is lined by trees on all four sides. (b) Depicting the arrangement of sampling points within the experimental field (blue dots). Values on the X and Y-axes represent arbitrary coordinates within a 27 × 27 matrix with coordinate (0,0) representing the centre. The dotted lines radiating out from the centre represent 16 compass directions corresponding to slices where wind directions were obtained (cardinal directions are not specified on this diagram). The red polygon represents the outline of the 21 × 21 matrix that was measured in the field (see red polygon in c). The position of the centre is off-set to increase sampling density in the direction of prevailing winds (see text for more information). (c) Aerial image showing the outline of the experimental field. Each red line along the square depicts the outer trap lines of each side of the sampling grid. The black and red place markers along the red diagonal represent the central release location (e.g. coordinate 0,0) and the weather station that was used to record weather parameters every 15 minutes respectively. (d) Diagram of the traps installed at sample locations. Fly not drawn to scale. See text for more detailed description.

The experimental design was constructed with sampling points positioned in a gradient radiating away from a central site from which the flies were released. The arrangement of each point was determined by first specifying a 21 × 21 matrix with 441 points (corresponding to 100 × 100 metre square with each point 5 m apart) and then by selecting specific points at which sampling locations were to be placed. Locations were selected by first converting Cartesian coordinates to polar coordinates using a trigonometric function that varied sampling concentration according to a cosine function and truncating the continuous values such that positive values represented sampling locations and negative values represented locations without sampling. This resulted in a total of 227 locations (including the central release location) that varied in distance and density in a parameterized gradient (Fig. 1b). Arranging sampling locations in this manner was done to balance the spatial resolution with the amount of time required to sample the entire field (i.e. temporal resolution). The experimental arena was prepared by first positioning each of the four corners 100 m apart in the field and then building trap lines within the arena. The arena was 100 × 100 m^2^ with 21 rows and 21 columns. Trap lines were positioned every 5 m. Stake flags (6.35 × 8.89 cm on a 53 cm wire stem, Milwaukee Tools) were placed at 5 m intervals along each trap line resulting in 441 flagged locations. The field diagram (Fig. 1b) was used to identify flags where a given sampling location was to be placed. One of the four corners of the arena was angled towards a North-Easterly direction. This was due to the prevailing winds mainly moving in an East and East-North-East direction as measured by our weather station (Vantage Vue, Davis Instruments, California, USA). For the same reason, the release site at coordinate (0,0) was off-set from the middle to allow higher sampling resolution in the direction of the prevailing winds. The weather station was installed along the diagonal (Fig. 1c) to measure wind direction and speed and temperature every 15 minutes.

Sampling locations consisted of baited traps inserted into the ground (10 cm off ground). Traps were prepared from a 90 mL plastic cup (Starplex, Starplex Scientific, Toronto, ON, Canada) positioned horizontally on a small piece of bamboo (Figure 1d). Prior to inserting a trap, the sampling location was prepared by manually removing vegetation around the flag and placing a small piece of landscaping fabric (The Scotts Company LLC) on the ground surface. To prevent other insects such as ants from gaining access to a trap, the small piece of bamboo that held the trap was inserted through the inner part of a two-part plastic shot glass (81.3 mL Amscan, Toronto, ON, Canada). The outer part of the glass was filled with water, which served as a moat and effectively prevented access to the piece of bamboo holding the baited trap (Fig. 1d).

Each trap was baited with 20 mL of fly medium consisting of a mixture of sugar, dead yeast and agar. Briefly, for approximately 1 L of medium we mixed 100 g of sugar, of which 50% of the sugar came from sucrose and the other 50 % came from bananas (~0.12 g of sucrose in 1 g of banana) that were blended prior to mixing, with 110g of yeast and 17.43 g of agar in 1 L of tap water. We also added 8 g of C_4_H_4_KNaO_6_, 1 g of KH_2_PO_4_ and 0.5 g each of NaCl, MgCl_2_, CaCl_2_ and Fe_2_(SO_4_)_3_ which was part of the standard yeast-sugar-agar laboratory medium that we used for maintaining fly stocks (Belay et al. 2007). All compounds were combined, mixed for an hour and autoclaved to ensure that the medium was sterile before being applied to traps. In a preliminary study in the laboratory, banana-baited traps were not biased towards any strain or sex used in the present study (results not shown).

### 2.3 Counting and marking flies

Each fly strain was counted and marked with a unique fluorescent pigment (see Edelsparre et al. 2014 for a full description of the marking technique). To estimate the number of flies we created a standard curve for the relationship between the number of flies and weight for each strain separately. This was done by counting and weighing batches of 50, 100, 150, 200, and 450 flies and estimating the linear regression equation for each strain. Each equation was used to predict the number of flies in each vial given their group weight and strain ID. Batches of approximately 150 flies were anesthetized with CO_2_ and transferred to separate empty plastic vials after which a minute amount of dry fluorescent pigment (DayGlo, Cleveland, OH, USA) was added (rover, Saturn Yellow, AX-17-N; sitter, Aurora Pink, AX-5-11; outbred flies, Horizon Blue, A-19). The colours were assigned randomly to each strain and were not known to the experimenters. Each vial was gently shaken to ensure that all flies were rolled in the pigment and subsequently transferred to 170 mL sponge-topped plastic *Drosophila* bottles where they were allowed 24 hours to groom themselves. This method left a badge on the ventral and dorsal thoraces that can be visualized by a portable black light in the field. In total 5644 rovers, 5352 sitters and 5657 outbred flies (2-7 days post eclosion) were tagged and released at the centre of the experimental field.

### 2.4 Releasing and recapturing flies

Prior to release, flies were acclimatized to their surroundings by placing the *Drosophila* bottles next to the central release trap at 09:00 on 10 October 2015. At 12:00 flies were released by gently removing the sponge-tops from each bottle in a haphazardly chosen order. It took 8 minutes from the release of the first bottle to the release of the last bottle.

Recapturing flies involved first randomly choosing between rows and columns of the trapping arena (Fig. 1b). Once either rows or columns were selected, three field observers sampled rows or columns in a random order. A complete sampling round involved visiting all 227 baited traps and bringing traps containing captured flies to a central location where all the samples were processed. Processing involved transferring flies to transparent *Drosophila* vials, identifying the colour markings by exposing flies to a black light and visually counting the number of flies with each colour at each sample location. After processing, the flies were transferred back to their respective baited trap and released at their capture location. Traps were baited with fresh food every second day. The first sampling round commenced 30 minutes after release (12:30), and we strove to complete a minimum of two sampling rounds per day over a course of five days (or until no or few flies remained on the experimental field). A sampling round took a maximum of two hours depending on how many flies were captured. In total 12 sampling rounds were completed over the 5 days, however, because of low capture rates after 50 hours post-release we used the first six sampling rounds in our analysis corresponding to 0.5, 1.5, 21, 26, 46 and 50 hours after the release (Fig. 2).

**Figure 2:**
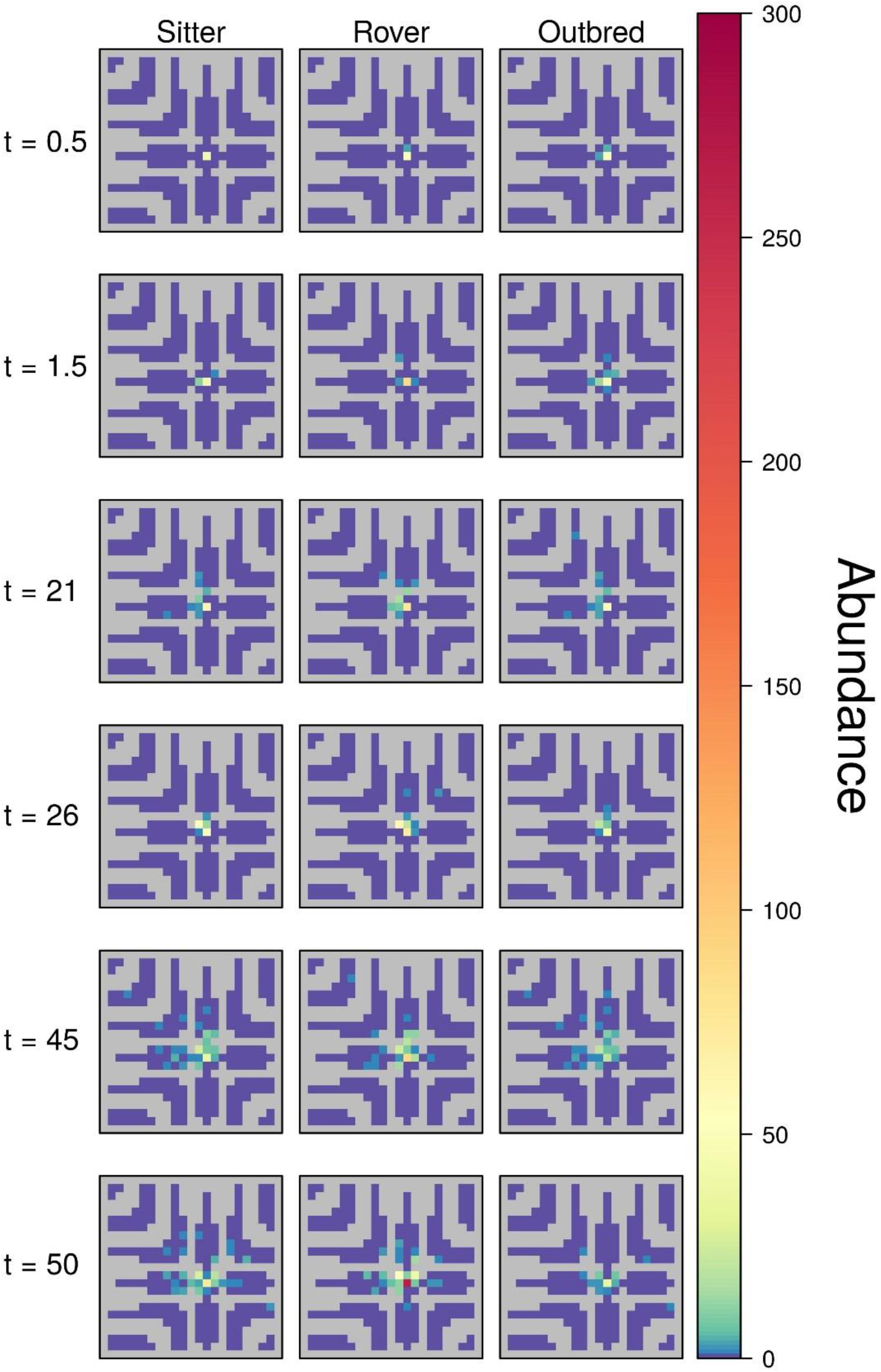
Starting from the top of the panel to the bottom we show time series of fly captures across the experimental field. Each square represents a time unique sequence. t=0.5, t=1.5, t=21, t=26, t=45 and t=50 represent captures at 30 minutes, 1.5 hr, 21 hrs, 26 hrs, 45 hrs and 50 hrs after release of the flies respectively. The left set of panels represent the time series for sitter (*for*^s^) flies and the middle and right set of panels represent the time series for rover (*for*^R^) and outbred flies respectively. The abundance of flies at each sample location are indicated by unique colour coding. Dark blue areas, as indicated by the draped legend on the right, indicate samples with zero fly captures and areas with darkest red indicate samples with abundances of maximum 300 flies. The gray area within each square indicate locations that did not contain traps.

### 2.5 Data analysis

To investigate how the rover, sitter and outbred strains respond to temperature and wind in the field, we developed and fit a dynamic spatio-temporal model to the data shown in Fig. 2. Dynamic spatio-temporal modelling enables mathematical models, like partial differential equations, to be fit to data using commonly applied parameter estimation techniques (Wikle et al. 2019; Hooten and Hefley 2019).

For each fly-strain, we fit a partial differential equation that included a component that describes ecological diffusion and a component that describes advection. By fitting a model to each strain separately we are able to evaluate how individual-level predictors (i.e. rover/sitter differences) translate to population-level processes. The partial differential equation can be written as

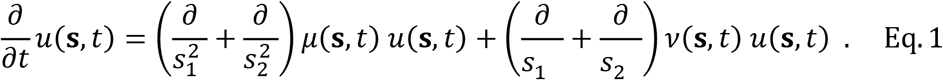

In Eq. 1, *u*(**s**, *t*) is the likelihood an individual is at location **s** ≡ (*s*_1_, *s*_2_)′ at time *t*, which can be converted to the intensity of the dispersing population by multiplying *θ* × *u*(**s**, *t*), where *θ* is the number of individuals released (i.e., 5644 rovers, 5352 sitters and 5657 outbred flies). The diffusion rate, *ν*(**s**, *t*), and advection rate, *μ*(**s**, *t*), can vary over both space and time and depend on environmental covariates (e.g., temperature and wind).

For the analysis, we specified the diffusion rate as follows

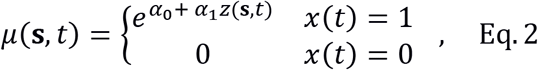

where *α*_0_ is the intercept and *α*_1_ is a regression coefficient for standardized temperature, *z*(**s**, *t*), at location **s** and time *t*. The indicator variable *x*(*t*) depends on the time, *t*, and is equal to 1 if it is daylight and equal to zero if it is night. When it is night we assume that the flies do not move (Konopka and Benzer 1971), thus the diffusion rate should be equal to zero. When it is daylight, the diffusion rate should always be greater than zero, thus motivating the exponential function used in Eq. 2. The standardized temperature was calculated by subtracting the mean temperature from the observed temperature and dividing this by standard deviation. Thus, the intercept term, *α*_0_, represents the natural log of the diffusion rate at the average temperature recorded during the study. Finally, temperature was measured at a single location, thus *z*(**s**, *t*) is spatially constant (i.e., *z*(**s**, *t*) ≡ *z*(*t*).

Similarly, we specified the advection rate as

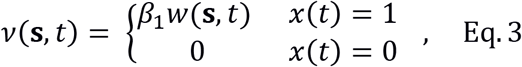

where *β*_1_ is a regression coefficient for wind velocity, *w*(**s**, *t*), at location **s** and time *t*. During daylight, the advection rate, *v*(**s**, *t*), can be positive or negative. Thus, there is no constraint required like the exponential function in Eq. 2. If the wind transports (advects) individuals, we expect a positive advection rate, whereas if individuals actively move into the wind then we expect a negative advection rate. Unlike the diffusion rate, the advection rate does not contain an intercept term because when the wind velocity is zero (i.e., *w*(**s**, *t*) = 0)), there should be no advection. Similar to temperature, wind velocity was measured at a single location, thus *w*(**s**, *t*) is spatially constant.

By fitting Eq. 1 to the data for each strain, we are able to estimate parameters that describe each strain’s movement in response to changes in the environment. For example, the magnitude and sign of the estimated value of *α*_1_ allows us to infer how the diffusion rate for each strain changes when the temperature increases. We provide a brief description below of how we fit partial differential equations to data. A more detailed description is provided in supporting material along with computer code (see Appendix S1).

Fitting Eq. 1 to the data from each strain first involves specifying a statistical model for the observed data. We assumed

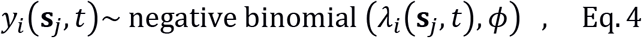

where *y*(**s**_*j*_, *t*) is the number of individual of strain *i* (*i* = 1,2,3) captured at the location **s**_*j*_ corresponding to the trap locations (i.e., *j* = 1,2,…,227) at the time *t*. For our data, the observed times were *t* = 0.5, 1.5,21,26,45, and 50 hours post release (see Fig. 2). For a given trap location and sampling time, the expected number of individuals caught from each strain was *λ*_*i*_(**s**_*j*_, *t*) which was modeled with

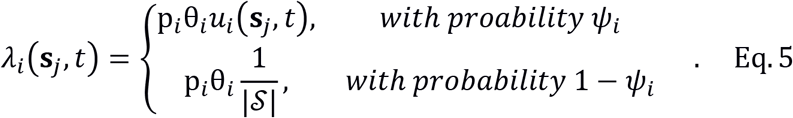

In Eq. 5, p_*i*_ is the capture probability, θ_*i*_*u*_*i*_(**s**_*j*_, *t*) is the intensity of the dispersing population from the Eq. 1 (where θ_*i*_ is the known number of individuals of strain *i* released), 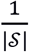 is a uniform probability density where 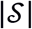 is the area of my study regions, and *ψ*_*i*_ is a mixture probability. Conceptually, Eq. 5 represents a population where, upon release, a proportion, *ψ*_*i*_, of the individuals’ movement is described by Eq. 1 and the movement of the remaining proportion of individuals (1 − *ψ*_*i*_) is described by a uniform distribution over the study area. The proportion of individuals with movement that follow the uniform distribution exhibit abnormal behaviour and could be caught with equal chances at any location within the study area. For example, upon release a small proportion of the flies exhibited bolting behaviour. Because these behaviours are abnormal, we expected that the estimated value of the parameter *ψ*_*i*_ would be close to one.

We took a Bayesian approach for parameter estimation and specified priors for all unknown parameters. To estimate the parameters in our model from the data, we developed software using a Markov chain Monte Carlo algorithm similar to the approach described in Hooten and Hefley (2019, Ch 28 pg. 501); however, solving Eq. 1 involved developing an emulator described by Hooten et al. (2011). Details of the implementation are provided in Appendix S1.

## 3. Results

In total, we captured 1223 rovers, 861 sitters and 279 outbred flies over the five days of the experiment. Approximately 95% (2363 flies) of all captured flies were captured within the first 50 hours of the experiment, corresponding to the first six sampling rounds. The capture data from the six sampling rounds used in the analysis are described in Figure 2. During the first 50 hours from release, the flies experienced a range of changing weather conditions, ranging from 10-22 °C during the day and nightly temperatures ranging from just above 5 °C to 16 °C (Fig. 3a). Wind speeds ranged from 0 to just above 2.5 m/s (Fig. 3b) with wind direction predominantly between 120 and 140 degrees, corresponding to East-North-East and East-South-East directions (Fig. 3c).

**Figure 3:**
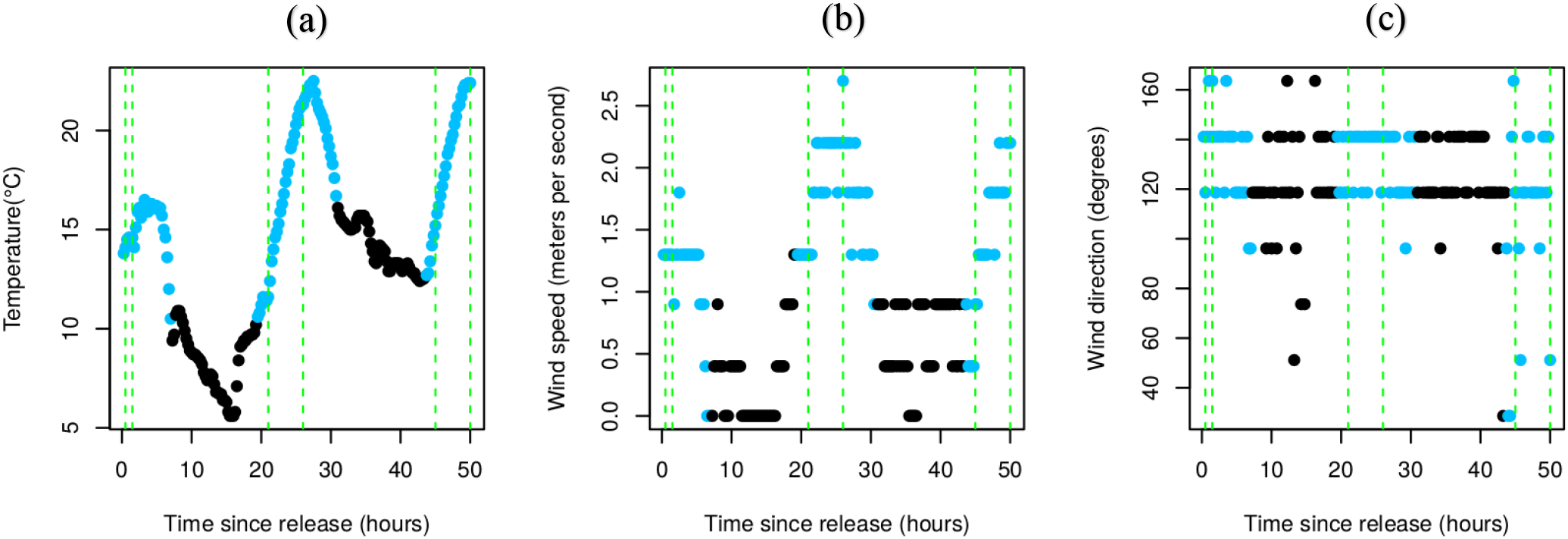
Temperature (a) and wind speed (b) and wind direction (c) measured every 15 minute (y-axes) over the 50 hours after release (x-axes). Each dot represents a temperature/wind speed/wind direction measurement, blue dots were measured during daylight and black dots represent measurements after daylight. The green vertical dashed lines in (a) (b) and (c) represent the temperature and wind measurements during the six sampling time points.

In general, our prediction that dispersal increased with temperature was supported. This conclusion is based on the finding that the posterior mean of the diffusion rates for rovers and sitters increased rapidly from 18 °C to 22 °C (Fig. 4). The posterior distribution of the diffusion rates show that dispersal of both sitters and rovers responded positively to temperature, but the magnitude of the response is uncertain, particularly at temperatures above 20 °C. In addition, the posterior mean estimates for rovers and sitters mirrored each other particularly at temperatures below 18 °C, but tended to be greater than the diffusion rate for outbred flies, which largely remained unchanged across the temperature range (Fig. 4). For example, if the temperature was 22 °C over a course of 24 hours, our results indicate that we should expect rovers and sitters to spread 79 m^2^ and 152 m^2^ more when compared to the outbred strain. Conversely, at temperatures below 14 °C the outbred strain spreads at least 36 m^2^ more over 24 hours when compared to rovers and sitters. The posterior distribution for all parameter estimates for the temperature analysis are shown in Fig. 6a and 6b.

**Figure 4:**
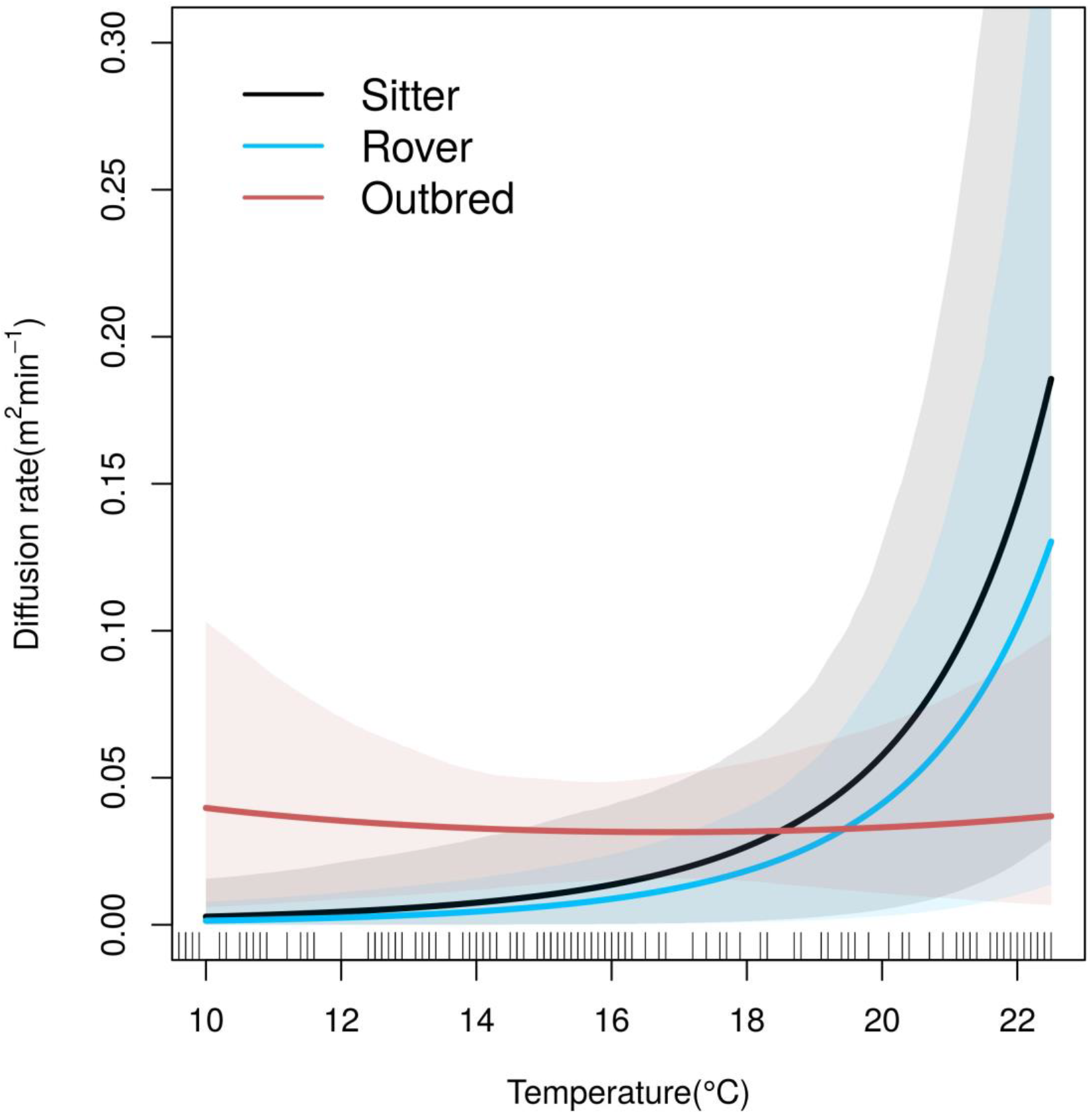
The relationship between temperature (x-axis) and the rate of movement (y-axis) for each fly strain. Each fly strain is represented by a bold line (blue = rovers, *for*^R^, black = sitters, *for*^s^ and red = outbred flies). The correspondingly coloured shaded areas surrounding each bold line represents the 95 % credible intervals for each fly strain. The thin vertical lines at the bottom of the x-axis indicate the range of temperatures that were measured during the course of this field experiment. At cooler temperatures the outbred strain tended to move faster than the rover and sitter strains, which showed very little movement. At warmer temperatures the diffusion rate of outbred flies remain largely unchanged while rover and sitter diffusion increased (i.e. at temperatures > 18 °C).

Wind speed and direction played a significant role in dispersal for all three fly strains. This conclusion is supported by the finding that the advection parameter was positive and increased with wind speed for rovers, sitters and outbred flies (Fig. 5). In fact, even with the level of uncertainty around each posterior mean the lower limit of the 95 % credible interval for all three strains remains above zero even at wind speeds below 0.5 m/s. Although there is uncertainty in the estimates (cf. 95 % credible intervals on Fig. 5), the difference among posterior mean responses becomes larger as wind speed increases. For example, sitters were most sensitive to wind speeds whereas rovers were the least sensitive; at a wind velocity of 1 m/s sitters are expected to advect approximately 5 metres further over 24 hours than rovers, however, at wind velocities of 2.5 m/s the difference between the two strains is expected to be approximately 13 metres over 24 hours. The parameter estimates for the advection part of the analysis are shown in Fig. 6c.

**Figure 5:**
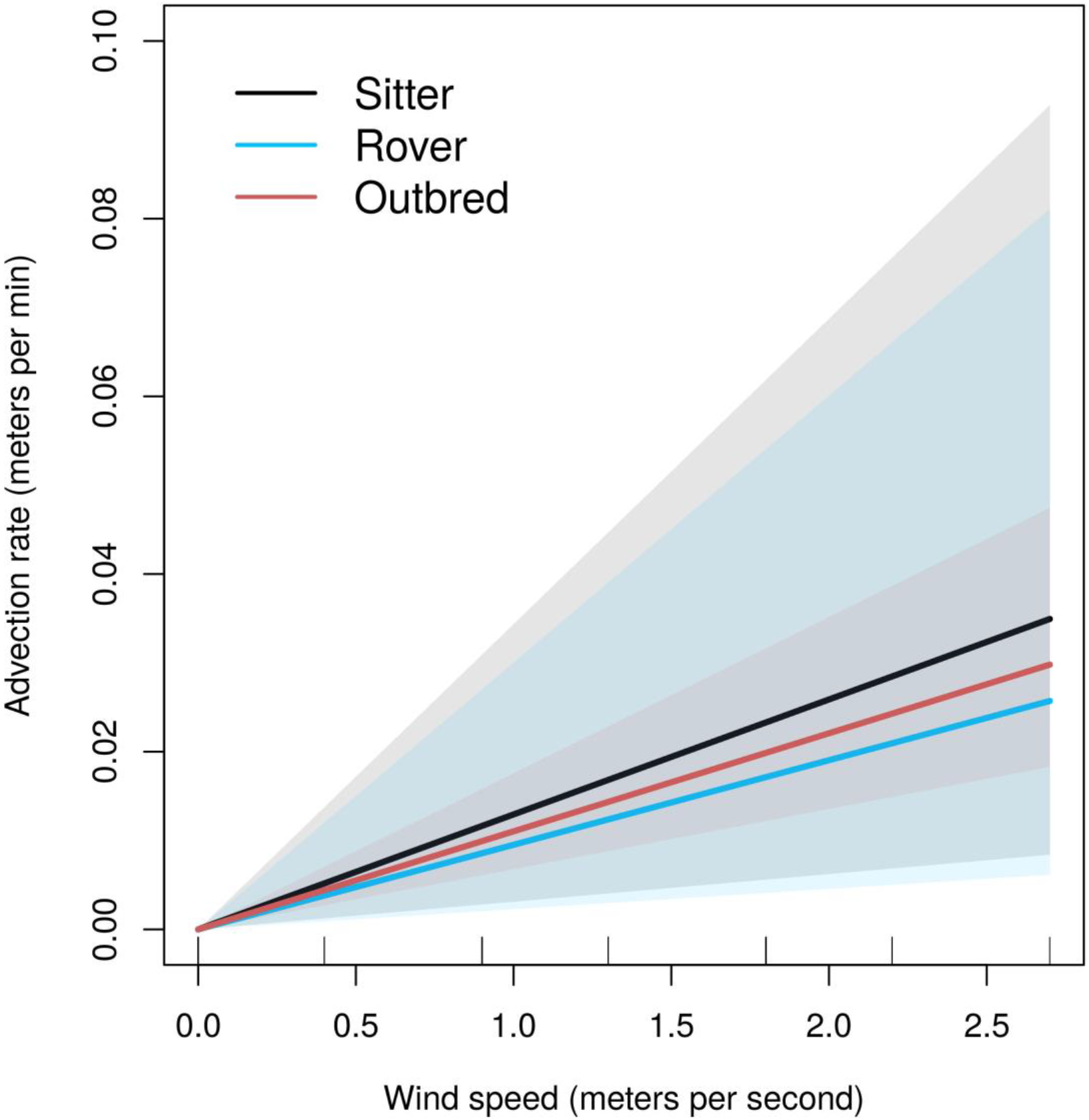
Depicting the relationship between wind speed (x-axis) and advection rate (y-axis). The advection rate (metres per minute) is positive if flies are dispersing in the direction of wind and negative if flies are dispersing against wind direction. For each fly strain the posterior means are represented by bold lines (blue = rovers, *for*^R^, black = sitters, *for*^s^ and red = outbred flies). The correspondingly coloured shaded areas surrounding each bold line represents the 95 % credible intervals for each fly strain. For all three strains the advection rate is positive and increases with wind speed.

**Figure 6:**
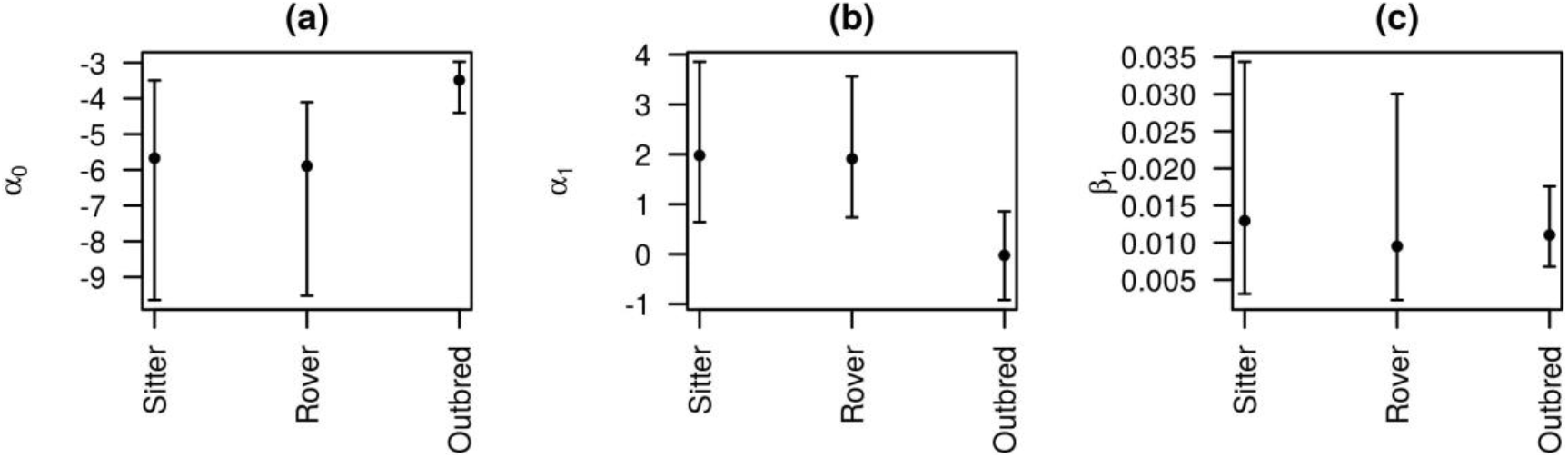
Parameter estimates for each fly strain associated with diffusion rate (Eq. 2, panels a and b), advection rate (Eq. 3, panel c. Each black dot represents the posterior mean with 95 % credible interval (vertical whiskers). In panel c positive values indicate the tendency to disperse with wind and negative values indicate the tendency to move against wind direction. The advection parameter is positive for all three fly strains.

## 4. Discussion

A number of conclusions can be drawn from the results of our study. First, temperature plays a critical role in fly dispersal in the field. We found that for rovers and sitters, the posterior mean of the diffusion rates increased with temperature particularly between 18 and 22 °C. In contrast, the posterior mean of the diffusion rate of the outbred fly strain tended to be greater than the diffusion rates of rovers and sitters at temperatures less than 18 °C, but largely remained unchanged across the temperature range. The rover and sitter responses to temperature mirror each other. However, the effect size of the mean estimates particularly between the rover and the outbred strain is noteworthy. For example, at 22 °C the mean diffusion rate is nearly four times larger for rovers than for the outbred strain (Fig. 4), suggesting that the models have the capacity to predict strain-dependent differences in dispersal outcomes over time. Second, wind plays a critical role in the dispersal of flies. The advection rate increased linearly with wind speed for all strains, however, rovers tended to be less sensitive to the effect of wind speed relative to sitters and outbred flies. As was the case for temperature, there is a relatively large amount of uncertainty in the magnitude of the response of sitters, rovers and outbred flies to wind speed, as indicated by the 95 % credible intervals, however, there is strong evidence that all three strains responded positively to increasing wind speed. In addition, the posterior mean advection distance between rovers and sitters more than doubled for every 1 m/s increase in wind speed (Fig. 5). This suggests that flies were influenced by both the direction and the speed of wind during the experiment, but that the magnitude of this effect tended to depend on fly-strain. This suggests that differences in down wind dispersal were due to behavioural differences and not a passive response. Thirdly, combining ecological diffusion modelling with wind advection clearly improved our ability to predict dispersal at the population-level. This conclusion is based on the finding that dispersal of flies in the field varied with both temperature and wind. For temperature, the posterior means of the diffusion rates increased rapidly for rovers and sitters, while the rate remained unchanged for the outbred population. For wind the effect is even stronger. The posterior means of the advection rates were above zero across the range of wind speeds for all three strains. Even with the level of uncertainty in the estimates, the lower limit of the 95 % credible intervals for all three posterior means is above zero at low wind speeds. This further strengthens the evidence in favour of a strong wind effect. Although there was some evidence of differences in response to wind speed, overall rovers, sitters and the outbred population exhibited similar patterns of response to wind (parallel responses). Taken together the environmental effects were strong for both the rover and sitter strains as well as the outbred population, although the strength of the evidence varied across the temperature and wind ranges. Our models therefore captured the effects of key descriptors of fly movement that a model without temperature and wind likely would have ignored and therefore our results inform our understanding of potential factors underlying dispersal in nature.

Climate driven environmental conditions such as temperature and wind have long been considered key factors influencing fly dispersal in particular and insect dispersal in general (Glick 1941, McManus 1988). Since the 1950’s prognoses of the relationship between temperature, wind, and dispersal have been used to both forecast and backtrack incidents of migrant pests such as African armyworms and desert locusts (Wellington 1954, Rainey 1979). In support of this, we found that wind speed and direction played a significant role in dispersal even at extremely low wind speeds. This suggests that wind transport can underlie a large part of the estimated rate of dispersal. We detected wind speeds between 0 and 3 m/s during this study. As such, wind may play an even larger role when speeds exceed 3 m/s. Clearly, in our study flies were able to navigate towards, around and into the traps. Taylor (1974) used the term “boundary layer” to describe a hypothetical layer of air near the ground where wind is not able to affect the movement of small insects (i.e. insect flight speed exceeds wind speed). Such a layer would allow flies to navigate towards and around traps. In our study, the wind above this boundary layer may have facilitated the dispersal of flies prior to visiting a particular trap, but whether or not the wind effect was mediated by behaviour (as a means to conserve energy) or remained passive cannot be directly answered by our data. Nevertheless, the low wind speeds and moderate to warm temperatures experienced during our study were conducive to insect movement (Glick 1942, Cockbain 1961, Taylor 1963).

Our findings demonstrate the potential use of genetic and environmental information in detecting individuals that may influence dispersal disproportionally. Although we did not address genetic effects directly in this study, we used different strains of flies, rovers (more dispersive) and sitters (less dispersive) and an outbred strain, to understand how flies with different genetic predispositions to disperse influenced the spatial spread of a fly-population. Unlike previous findings in the laboratory and the field (Edelsparre et al. 2014, Edelsparre et al. 2018), the known difference in dispersal between rovers and sitters was minimized in response to temperature and wind conditions. Sitters tended to disperse faster with wind than did rovers particularly at high wind speeds. This suggests that an understanding of dispersal likely hinges on disentangling how genes involved in dispersal interact with relevant factors in the environment to affect individual differences in dispersal behaviour (Sokolowski 2001, Dudaniec et al. 2018, Saastamoinen et al. 2018). Insect movement involves multiple genes (Saastamoinen et al. 2018) in interaction with many environmental factors including temperature and wind measured in the present study. *for* is one of the genes that affects movement related behaviours in a wide range of insect species, including ants (Ingram et al. 2005) and honey bees (Ben-Shahar et al. 2002). *for* has also recently been associated with outbreaks of locusts (Tobbak et al. 2013) and possibly spruce budworm, *Choristoneura fumiferana* (Van Hezewijk et al. 2018). Describing the role of *for* and other candidate genes in invasive species should provide a unique opportunity to better understand and model invasion biology in terrestrial environments. Additionally, by extending this thinking to include temperature and wind our findings are particularly pertinent to elucidating conditions under which climatic factors may influence insect invasions. Specific predictions regarding climatic effects on insect dispersal offer the potential to reduce uncertainty in invasion biology where unpredictability seems to be the rule rather than the exception (Melbourne and Hastings 2009).

Our findings provide a powerful framework for combining individual-level predictors with climate driven variables to understand dispersal at the population-level. An implicit assumption in many models, including diffusion models, is that the dispersing individuals within the population are identical (Kot et al. 1996, Gurarrie et al. 2009). Other studies have proposed individual differences in behaviour as an explanation for non-random variation in dispersal frequently reported in the literature (Skalski and Gilliam 2000, Fraser et al. 2001, Réale et al. 2007). We addressed this assumption directly by using the rover, sitter and outbred strains of flies as proxies for individual differences in dispersal propensity. Through the fitting of separate models for each strain we show how such efforts can be useful predictors of population-level dispersal in the field. Our results are consistent with models that propose individual variation in behaviour as explanations for population-level variation in dispersal. Our findings not only provide the potential to inform how individuals influence dispersal disproportionally but they also improve our understanding of key mechanisms surrounding individual differences in response to environmental factors that give rise to population-level dispersal.

There are potential limitations to our study. First, the temporal resolution we used in our experimental design affected the power of our analysis. Increasing temporal sampling over spatial sampling could have provided stronger evidence for differences among strains. One way to increase the temporal resolution may be to sample flies as presence/absence data rather than abundance. This would reduce the time spent at each sampling location and increase the number of times the entire arena could be surveyed in a day. Second, the rover and sitter strains have been cultured in the laboratory for over 30 years and may not have recapitulated dispersal behaviour in natural populations. The dispersal rate of rovers and sitters above 18 °C showed a response to temperature consistent with insect flight activity reported in the literature (Cockbain 1961, Taylor 1963). Along with the rover and sitter strains, we released an outbred strain established from the field. This strain was assembled in 2015 using 92 isofemale lines collected in 2012. The diffusion rate of this outbred strain was stable across the temperature range in comparison to the rover and sitter strains. The outbred strain was collected near the limit of *D. melanogaster’s* northern-most distribution in Sudbury Ontario, Canada (Thomas Merritt, personal communication). This might explain why the outbred strain tended to be more active at low temperatures than the rover and sitter strains that were originally caught well within *D. melanogaster’s* normal range (Toronto, Ontario, Canada, 402 km south of Sudbury). Overall, our results provide a useful approach for investigating how behavioural differences might lead to dispersal outcomes for a population with multiple dispersal strategies and how those strategies may interact with temperature and wind.

Our study provides two scientific contributions to our understanding of the evolution and ecology of organismal movement. First, we took advantage of the wealth of information available for flies in general and this *Drosophila melanogaster* model system in particular to build models that incorporated genetic variation known to underlie dispersal strategy (Edelsparre et al. 2014, Edelsparre et al. 2018) and climatic effects. Earlier attempts to understand the movement of organisms generally relied on simpler analyses of linear rates of spread involving individuals with equal dispersal propensity (Hastings 2005, Cote et al. 2017). A long history of publications rooted in the ecology of organismal movement illuminate a far more complex and interesting process and over the last two decades researchers have pushed for an integration of behavioural and environmental data with movement theories expressed as mathematical models (Hastings et al. 2005, Nathan 2008, Hefley et al. 2017). In line with this idea, our results clearly demonstrate that more complex models indeed can be critical to capturing factors that influence animal movement at the individual level (i.e. multiple dispersal strategies) and highlight the importance of matching theoretical models of movement with data to better understand the distribution of organisms in nature. Second, we developed a powerful framework for linking individual variation in dispersal strategy with the environment. Understanding how such components can lead to population-level dispersal is valuable because individual differences in behaviour are increasingly linked with important ecological and evolutionary processes such as habitat fragmentation (Cote et al. 2017), climate change (Fitzpatrick and Edelsparre 2018), and biogeography (Canestrelli et al. 2016). In fact, individual variation in behaviour has recently been referred to as a ‘pacemaker’ of evolution of non-behavioural traits. Our study therefore offers a unique opportunity to understand how dispersal strategy influences the distribution of populations and ultimately the pattern of biological diversity on our planet (Canestrelli et al. 2016).

## Acknowledgements

We acknowledge that this study was carried out on traditional and sovereign territory of the Onkwehonwe Peoples of Six Nations of the Grand River and the Anishinaabe Peoples of the Mississaugas of the New Credit and is within the territory of the Neutral Peoples. This study was possible because of the historical and ongoing indigenous stewardship of these lands and remain critical to biodiversity in Ontario. We also thank ***rare*** Charitable Research Reserve, Cambridge, Ontario for facilitating our use of the field site, providing field equipment as well as providing access to field laboratory and accommodation during our stay at the reserve. We would particularly like to thank Jenna Quinn the Program Scientist at ***rare*** for making our stay at ***rare*** smooth and enjoyable. We are deeply grateful for the assistance of Anders Vesterberg and Jenna Chen whose clever and meticulous efforts helped setting up and executing the experiment as well as collecting the data. Lastly, we are grateful to Thomas Merritt for collecting the iso-female lines that produced the outbred population. This research was funded by Natural Sciences and Engineering Research Council (NSERC) to MJF, MAR and MBS. Partial funding for this project was provided by the National Science Foundation (NSF) via grant DEB 1754491 to TJH.

## References

Anreiter, I., and M. B. Sokolowski. 2019. The *foraging* gene and its behavioral effects: pleiotropy and plasticity. Annual Review Genetics. 53: 373–392.

Belay, A. T., R. Scheiner, A. K.-C. So, S. J. Douglas, M. Chakaborty-Chatterjee, J. D. Levine and M. B. Sokolowski. 2007. The *foraging* gene of *Drosophila melanogaster*: spatial-expression analysis and sucrose responsiveness. Journal of Comparative Neurology. 504: 570–582.

Ben-Shahar Y., A. Robichon, M. B. Sokolowski, and G. E. Robinson. 2002. Influence of gene action across different time scales on behavior. Science. 296: 741–744.

Brisson, J. A. 2010. Aphid wing dimorphisms: linking environmental and genetic control of trait variation. Philosophical Transactions of the Royal Society B. 365: 605–616.

Bocxlaer, I. V., S. P. Loader, K. Roelants, M. Menegon and F. Bossuyt. 2010. Gradual adaptation toward a range-expansion phenotype initiated the global radiation of toads. Science. 327: 679–682.

Canetrelli, D., R. Bisconti and C. Carere. 2016. Bolder takes all? The behavioral dimension of biogeography. Trends in Ecology and Evolution. 31: 35–43.

Clarke, J. S., S R. Carpenter, M. Barber, S. Collons, A. Dobson, J. A. Foley, D. M. Lodge, M. Pascual, R. Pielke Jr., W. Pizer, C. Pringle, W. V. Ried, K. A. Rose, O. Sala, W. H. Schlesinger, D. H. Wall and D. Wear. 2001. Ecological forecasts: an emerging imperative. Science. 293: 657–660.

Cote, J. and J. Clobert. 2007. Social personalities influence natal dispersal in a lizard. Proceedings of the Royal Society B. 274: 383–390.

Cote, J., E. Bestion, S. Jacob, J. Travis, D. Legrand and M. Baguette. 2017. Evolution of dispersal syndromes in fragmented landscapes. Ecography. 40: 56–73.

Cockbain, A. J. 1961. Low temperature thresholds for flight in *Aphis fabae* Scop. Entomologia Experimentalis et Applicata. 4: 211–219.

Coyne, J. A., I. A. Boussy, T. Prout, S. H. Bryant, J. S. Jones and J. A. Moore. 1982. Long-distance migration of *Drosophila*. The American Naturalist. 119: 589–595.

Denno, R. F., Roderick, G. K., Olmstead, K. L., & Döbel, H. G. 1991. Density-related migration in planthoppers (*Homoptera: Delphacidae*): the role of habitat persistence. The American Naturalist. 138: 1513–1541.

Duckworth, R. A. and A. V. Badyaev. 2007. Coupling of dispersal and aggression facilitates the rapid range expansion of a passerine bird. Proceedings of the National Academy of Sciences USA. 104: 15017–15022.

Dudaniec, R. Y., C. J. Yong, L. T. Lancaster, E. I. Svensson, and B. Hansson. 2018. Signatures of local adaptation along environmental gradients in a range-expanding damselfly (*Ischnura elegans*). Molecular Ecology. 27: 2576–93.

Edelsparre, A. H., M. J. Fitzpatrick, M. A. Rodríguez and M. B. Sokolowski. 2020. Tracking dispersal across a patchy landscape reveals a dynamic interaction between genotype and habitat structure. Oikos. DOI: https://doi.org/10.1111/oik.07368.

Edelsparre, A.H., McLaughlin, R.L. & Rodríguez, M.A. 2013. Risk taking not foraging behavior predicts dispersal of recently emerged stream brook charr (*Salvelinus fontinalis*). Ecosphere. 4: 73.

Edelsparre, A. H., S. Shahid and M. J. Fitzpatrick. 2018. Habitat connectivity is determined by the scale of habitat loss and dispersal strategy. Ecology and Evolution. 8: 5508–5514.

Edelsparre, A. H., A. Vesterberg, J. H. Lim and M. J. Fitzpatrick. 2014. Alleles underlying larval foraging behaviour influence adult dispersal in nature. Ecology Letters. 17: 333–339.

Fitzpatrick, M. J. and A. H. Edelsparre. 2018. The genomics of climate change. Science. 359: 29–30.

Freedman, L. A., S. Conant and R. C. Fleischer. 1987. Evolutionary ecology and radiation of Hawaiian passerine birds. Trends in Ecology and Evolution. 2: 196–203.

Fraser, D. F., J. F. Gilliam, M. J. Daley, A. N. Le and G. T. Skalski. 2001. Explaining leptokurtic movement distributions: intrapopulation variation in boldness and exploration. The American Naturalist. 158: 124–135.

Garlick, M. J., J. A. Powell, M. B. Hooten and L. R. McFarlane. 2012. Homogenization of large-scale movement models in ecology. Bulletin of Mathematical Biology. 73: 2088–2108.

Glick, P. A. 1939. The distribution of insects, spiders, and mites in the air. United States Department of Agriculture Technical Bulletin. 673: 1–151.

Glick, P. A. 1942. Insect population and migration in the air. pp. 88–98 in Moulton, S. (Ed.), Aerobiology. American Association of Advanced Science. Washington, D.C. A.A.A.S. Publ. 17.

Gomez-Uchida, D., D. Canas-Rojas, C. M. Riva-Rossi, J. E. Ciancio, M. A. Pascual, B. Ernst, E. Aedo, S. S. Musleh, F. Valenzuela-Aguayo et al. 2018. Genetic signals of artificial and natural dispersal linked to colonization of South America by non-native Chinook salmon (*Oncorhynchus tshawytscha*). Ecology and Evolution. 00: 1–18.

Grant, P. R. 1981. Speciation and the adaptive radiation of Darwin’s finches. American Scientist. 69: 653–663.

Gurarie, E., J. J. Anderson and R. W. Zabel. 2009. Continuous models of population-level heterogeneity inform analysis of animal dispersal and migration. Ecology. 90: 2233–2242.

Hanski, I. and M. Gilpin. 1991. Metapopulation dynamics: brief history and conceptual domain. Biological Journal of the Linnean Society. 42: 3–16.

Hastings, A., K. Cuddington, K. F. Davies, C. J. Dugaw, S. Elmendorf, A. Freestone, S. Harrison, M. Holland, J. Lambrinos et al. 2005. The spatial spread of invasions: new developments in theory and evidence. Ecology Letters. 8: 91–101.

Hefley, T. J., M. B. Hooten, R. E. Russel, D. P. Walsh and J. A. Powell. 2017. When mechanism matters: Bayesian forecasting using models of ecological diffusion. Ecology Letters. 20: 640–650.

Hilbe, J. M. (2011). Negative Binomial Regression. Cambridge University Press.

Hill, J. K., C. D. Thomas, D. S. Blakeley. 1999. Evolution of flight morphology in a butterfly that has recently expanded its geographic range. Oekologia. 121: 165–170.

Hooten, M. B., M. J. Garlick, and J. A. Powell. 2013. Computationally efficient statistical differential equation modeling using homogenization. Journal of Agricultural, Biological, and Environmental Statistics. 18: 405–428.

Hooten, M. B., Leeds, W. B., Fiechter, J., & Wikle, C. K. (2011). Assessing first-order emulator inference for physical parameters in nonlinear mechanistic models. Journal of Agricultural, Biological, and Environmental Statistics. 16: 475–494.

Hooten, M.B. and T.J. Hefley. 2019. Bringing Bayesian Models to Life. Chapman and Hall/CRC.

Ingram K. K., P. Oefner, D. M. Gordon. 2005. Task-specific expression of the foraging gene in har-vester ants. Molecular Ecology. 14: 813–818.

Johnson, C. G. 1969. Migration and Dispersal of Insects by Flight. Methuen & Co Ltd. London. 763 pp.

Konopka, R. J. and S. Benzer. 1971. Clock mutants of *Drosophila melanogaster*. Proceedings of the National Academy of Sciences of the United States of America. 68: 2112–2116.

Korsten, P., T. van Overveld, F. Adriaensen and E. Matthysen. 2013. Genetic integration of local dispersal and exploratory behaviour in a wild bird. Nature Communications. 4: 2362.

Kot, M., M. A. Lewis and P. van den Driessche. 1996. Dispersal data and the spread of invading organisms. Ecology. 77: 2027–2042.

Lombaert, E., A. Estoup, B. Facon, B. Joubart, J. C. Grégoire, A. Jannin, A. Blin, and T. Guillemaud. 2014. Rapid increase in dispersal during range expansion in the invasive ladybird *Harmonia axyridis*. Journal of Evolutionary Biology. 27: 508–517.

Losos, J. B., T. R. Jackman, A. Larson, K. de Queiros and L. Rodríguez-Schettino. 1998. Contingency and determinism in replicated adaptive radiations of island lizards. Science. 279: 2115–2118.

Markow, T. A. and S. Castrezena. 2000. Dispersal in cactophilic *Drosophila*. Oikos. 89: 378–386.

Melbourne, B. A., and A. Hastings. 2009. Highly variable spread rates in replicated biological invasions: fundamental limits to predictability. Science. 325: 1536–1539.

McManus, M. L. 1988. Weather, behaviour and insect dispersal. Memoirs of the Entomological Society of Canada. 146: 71–94.

Melbourne, B. A. and A. Hastings. 2009. Highly variable spread rates in replicated biological invasions: fundamental limits to predictability. Science. 325: 1536–1539.

Myles-Gonzalez, E., G. Burness, S. Yavno, A. Rooke and M. G. Fox. 2015. To boldly go where no goby has gone before: boldness, dispersal tendency, and metabolism at the invasion front. Behavioural Ecology. 26: 1083–1090.

Nathan, R. 2008. An emerging movement ecology paradigm. Proceedings of the National Academy of Sciences of the United States of America. 105: 19050–19051.

Nathan, R., W. M. Getz, E. Revilla, M. Holyoak, R. Kadmon, D. Saltz and P. Smouse. 2008. A movement ecology paradigm for unifying organismal movement research. Proceedings of the National Academy of Sciences of the United States of America. 105: 19052–19059.

O’Riain, M. J., J. U. M. Jarvis and C. Faulkes. 1996. A dispersive morph in the naked mole rat. Nature. 380: 619–621.

Osborne, K. A., A Robichon, E. Burgess, S. Butland, R. A. Shaw, A. Coulthard, H. S. Pereira, R. J. Greenspan and M. B. Sokolowski. 1987. Natural behavior polymorphism due to a cGMP-dependent protein kinase of *Drosophila*. Science. 227: 834–836.

Phillips, B. L. 2009. The evolution of growth rates on an expanding range edge. Biology Letters. 5: 802–804.

Phillips, B. L., G. P. Brown, J. K. Webb and R. Shine. 2006. Invasion and the evolution of speed in toads. Nature. 439: 803.

Rainey, R. C. 1979. Interactions between weather systems and population of locusts and noctuids in Africa. pp. 109–119. In Rabb, R. L. and G. G. Kenedy (Eds.), Movement of highly mobile insects: concepts and methodology in research. North Carolina State Univ. Raleigh. 455 pp.

Réale, D., S. M. Reader, D. Sol, P. T. McDougall and N. J. Dingemanse. 2007. Integrating animal temperament with ecology and evolution. Biological Reviews. 82: 291–318.

Richardson, R. H. and J. S. Johnston. 1975. Behavioral components of dispersal in *Drosophila mimica*. Oecologia. 20: 287–299.

Ronce, O. 2007. How does it feel to be like a rolling stone? Ten questions about dispersal evolution. Annual Review of Ecology, Evolution and Systematics. 38: 231–253.

Saastamoinen, M., G. Bocedi, J. Cote, D. Legrand, F. Guillaume, C. W. Wheat, E. A. Fronhofer, C. Garcia, R. Henry, A. Husby, et al. 2018. Geneticsof dispersal. Biological Reviews. 93: 574–599.

Skalski, G. T. and J. F. Gilliam. 2000. Modeling diffusive spread in a heterogeneous population: a movement study with stream fish. Ecology. 81: 1685–1700.

Skellam, J. G. 1951. Random dispersal in theoretical populations. Biometrika. 38: 196–218.

Sokolowski, M. B. 2001. Genetics meets behaviour. Nature Reviews Genetics. 2: 879–890.

Sokolowski, M. B., C. Kent and J. Wong. 1984. *Drosophila* larval foraging behaviour: developmental stages. Animal Behaviour. 32: 645–651.

Taylor, L. R. 1963. Analysis of the effect of temperature on insects in flight. Journal of Animal Ecology. 32: 99–117.

Taylor, L. R. 1974. Insect migration, flight periodicity, and the boundary layer. Journal of Animal Ecology. 43: 225–238.

Tobback, J., H. Verlinden, K. Vuerinckx, R. Vleugels, J. Vanden Broeck & R. Huybrechts. 2013. Developmental-and food-dependent *foraging* transcript levels in the desert locust. Insect Science. 20: 679–688.

Van Hezewijk B., D. Vertman, D. Stewart, C. Béliveau, and C. Cusson. Environmental and genetic influences on the dispersal propensity of spruce budworm (*Choristoneura fumiferana*). 2018. Agricultural and Forest Entomology. 20: 433–441.

Wainwright, C. E., P M. Stepanian, D. R. Reynolds and A. M. Reynolds. 2017. The movement of small insects in the convective boundary layer: linking pattern with process. Scientific Reports. 7: 5438.

Wellington, W. G. 1954. Atmospheric circulation processes and insect ecology. The Canadian Entomologist. 86: 312–333.

Wikle, C. K. 2003. Hierarchical Bayesian models for predicting the spread of ecological processes. Ecology. 84: 1382–1394.

Wikle, C.K., A. Zammit-Mangion, and N. Cressie. 2019. Spatio-Temporal Statistics with R. Chapman and Hall/CRC.

Zera, A. J. and R. F. Denno. 1997. Physiology and ecology of dispersal polymorphism in insects. Annual Review of Entomology. 42: 207–230.

