## Appendix S1 for "Scaling up: understanding movement from individual differences to population-level dispersal"

### Appendix S1: Reproducible Analysis

### 1 Introduction

This appendix provides additional explanation and R code associated with data analysis presented in the manuscript. In addition, this appendix can be used to reproduce all results and figures presented in the manuscript. The R code in this appendix was designed to be executed in the order presented due to dependence on prior code. We begin by loading the required R packages.

```
library(sp)
library(raster)
library(rasterVis)
library(gridExtra)
library(grid)
library(matrixcalc)
library(gbm)
library(plotrix)
library(MASS)
```

### 2 Data

#### 2.1 Download data

Next we need to load the data. After publication, the data will be available on the Dryad Digital repository (Edelsparre et al. 2020). For the purpose of peer review, two urls are provided in the code below. The code below will automatically download the data and read it into R.

```
url <- "https://www.dropbox.com/s/th482j4bfkc6gv5/experimental%20data.csv?dl=1"
df.flies <- read.csv(url)

url <- "https://www.dropbox.com/s/mkxe8zr7p6hmc5a/weather%20data.csv?dl=1"
df.weather <- read.csv(url)
```

#### 2.2 Format data

Before the data are ready for analysis, we need to take several steps. The code below transforms the data into the format that was used for this example.

```
### Fly capture data

# Remove observations 50 hours after release because few individual were captured
df.flies <- df.flies[which(df.flies$Hours <= 50), ]
df.flies <- df.flies[which(df.flies$Hours > 0), ]

# 227 sample locations, 6 types (genotype x sex), 6 time points 227*6*6 = 8172 observations
dim(df.flies)

# Make point file of trap locations
pts.trap <- SpatialPointsDataFrame(df.flies[, 4:5], data.frame(capture = df.flies$Capture,
    time = df.flies$Hours, sex = df.flies$Sex, genotype = df.flies$Genotype))

# Make raster layer of study area and sampling grid
rl.capture <- raster(, xmn = -65, xmx = 50, ymn = -50, ymx = 65, res = 5, crs = NA)

# Associate fly counts with sampling grid cells
for (i in 1:length(unique(df.flies$Hours))) {
    t <- unique(df.flies$Hours)[i]
    temp <- rasterize(pts.trap[which(pts.trap$time == t & pts.trap$genotype == "Rover"), 1],
        rl.capture, field = "capture", fun = "sum")
    rl.capture.rover <- if (i == 1) {
        temp
    } else {
        stack(rl.capture.rover, temp)
    }
```

```
    temp <- rasterize(pts.trap[which(pts.trap$time == t & pts.trap$genotype == "Sitter"), 1],
        rl.capture, field = "capture", fun = "sum")
    rl.capture.sitter <- if (i == 1) {
        temp
    } else {
        stack(rl.capture.sitter, temp)
    }

    temp <- rasterize(pts.trap[which(pts.trap$time == t & pts.trap$genotype == "Sudbury"),
        1], rl.capture, field = "capture", fun = "sum")
    rl.capture.sudbury <- if (i == 1) {
        temp
    } else {
        stack(rl.capture.sudbury, temp)
    }
}
names(rl.capture.rover) <- paste("t=", unique(df.flies$Hours), sep = "")
names(rl.capture.sitter) <- paste("t=", unique(df.flies$Hours), sep = "")
names(rl.capture.sudbury) <- paste("t=", unique(df.flies$Hours), sep = "")

### Weather data

# Remove weather data that was recorded more than 50 hours after the release
df.weather <- df.weather[which(unique(df.weather$Hours) <= 50), ]

# Make raster layer for weather data.  Should be exactly the same dimensions and resolution
# as rl.capture
rl.weather <- raster(, xmn = -65, xmx = 50, ymn = -50, ymx = 65, res = 5, crs = NA)
rl.weather[] <- 0
rl.weather <- stack(mget(rep("rl.weather", length(df.weather$Hours))))
rl.tempout <- rl.weather
rl.wind_ew <- rl.weather
rl.wind_ns <- rl.weather

# Associate weather variables with sampling grid cells Positive values of
# cos((90-df.weather$relWindDir)/57.27) means flies will be advected from 'right' to 'left'
# Positive values of sin((90+df.weather$relWindDir)/57.27) means flies will be advected
# from 'top' to 'bottom'
for (t in 1:dim(rl.weather)[3]) {
    rl.tempout[[t]] <- df.weather$tempOut[which(df.weather$Hours == sort(df.weather$Hours)[t])]
    rl.wind_ew[[t]] <- ifelse(df.weather$WindSpeed == 0, 0, df.weather$WindSpeed * cos((90 -
        df.weather$relWindDir)/57.27))[which(df.weather$Hours == sort(df.weather$Hours)[t])]
    rl.wind_ns[[t]] <- ifelse(df.weather$WindSpeed == 0, 0, df.weather$WindSpeed * sin((90 +
        df.weather$relWindDir)/57.27))[which(df.weather$Hours == sort(df.weather$Hours)[t])]
}
```

#### 2.3 Figure 2

Below is R code that can be used to reproduce Figure 2.

```
# pdf(file='Fig_2.pdf',width=5,height=8,onefile=FALSE)
myTheme <- BTCTheme()
myTheme$panel.background$col = "gray"
rl.capture.all <- stack(rl.capture.sitter, rl.capture.rover, rl.capture.sudbury)
rl.capture.all <- crop(rl.capture.all, extent(c(-65, 50, -50, 65)))
levelplot(rl.capture.all, cuts = 254, at = c(seq(0, 300, by = 1)), margin = FALSE, layout = c(3,
    6), index.cond = list(c(1, 7, 13, 2, 8, 14, 3, 9, 15, 4, 10, 16, 5, 11, 17, 6, 12, 18)),
    scales = list(draw = FALSE), names.attr = c("Sitter", rep("", 5), "Rover", rep("", 5),
        "Outbred", rep("", 5)), col.regions = colorRampPalette(rev(brewer.pal(11, "Spectral")),
        bias = 2.5), par.settings = myTheme)
grid.text("Abundance", x = 0.94, y = 0.5, just = c("center", "bottom"), gp = gpar(cex = 1.7),
    rot = -90)
for (i in 1:length(unique(df.flies$Hours))) {
```

```
    grid.text(paste("t =", unique(df.flies$Hours))[i], x = 0.01, y = 0.88 - 0.1565 * (i - 1),
        just = c("left", "center"), gp = gpar(cex = 1), rot = 0)
}
# dev.off()
```

##### 2.4 Figure 3

Below is R code that can be used to reproduce Figure 3.

```
# pdf(file='Fig_3.pdf',width=8.5,height=3.3)
par(mfrow = c(1, 3))
plot(df.weather$Hours, df.weather$tempOut, col = c("black", "deepskyblue")[df.weather$Daylight +
    1], pch = 19, xlab = "Time since release (hours)", ylab = expression("Temperature(" * degree *
    C * ")"), main = "(a)")
abline(v = unique(df.flies$Hours), col = "green", lty = 2)
plot(df.weather$Hours, df.weather$WindSpeed, col = c("black", "deepskyblue")[df.weather$Daylight +
    1], pch = 19, xlab = "Time since release (hours)", ylab = "Wind speed (meters per second)",
    main = "(b)")
abline(v = unique(df.flies$Hours), col = "green", lty = 2)
plot(df.weather$Hours, df.weather$relWindDir, col = c("black", "deepskyblue")[df.weather$Daylight +
    1], pch = 19, xlab = "Time since release (hours)", ylab = "Wind direction (degrees)", main = "(c)")
abline(v = unique(df.flies$Hours), col = "green", lty = 2)
# dev.off()
```

#### 3 Statistical analysis

As described in the manuscript, we fit an advection-diffusion model to the experimental data. In what follows, we present the statistical model and algorithm used to implement it. Readers unfamiliar with mechanistic spatio-temporal statistical models will find introductions in Hooten and Hefley (2019; pgs. 501–515) and Wikle et al. (2019; ch. 5) useful. Equations that appear in this supporting material are labeled in the order they are presented, but a preceded by the letter S.

##### 3.1 Bayesian Model

For completeness, we re-present the advection-diffusion model and include some additional details. For our analysis we used

$$y_i(\mathbf{s}_j, t) \sim \text{NB}(\lambda_i(\mathbf{s}_j, t), \phi)\,, \tag{S1}$$

where $y_i(\mathbf{s}_j, t)$ is the number of individuals of strain $i$ ($i = 1, 2, 3$) captured at time $t$ and location $\mathbf{s}_j$. For our data, the observed times were $t = 0.5, 1.5, 21, 26, 45$ and 50 hours post release and there were 227 trap locations (i.e., $j = 1, 2, ..., 227$). The vector $\mathbf{s} \equiv (s_1, s_2)'$ contains the coordinates of a point within the study area (see Fig. 1). The abbreviation NB is a negative binomial distribution with $\text{E}(y_i(\mathbf{s}_j,t)) = \lambda_i(\mathbf{s}_j, t)$ and $\text{Var}(y_i(\mathbf{s}_j,t)) = \lambda_i(\mathbf{s}_j, t) + \phi\lambda_i(\mathbf{s}_j, t)$.

The expected number of individuals of a particular strain captured at time $t$ and trap location $\mathbf{s}_j$ is

$$\lambda_i(\mathbf{s}_j, t) = \begin{cases} p_i\theta_i u_i(\mathbf{s}_j, t) & \text{, with probability } \psi_i \\ p_i\theta_i \frac{1}{|\mathcal{S}|} & \text{, with probability 1-}\psi_i \end{cases}. \tag{S2}$$

In Eq. S2, $p_i$ is the unknown capture probability, $\theta_i$ is the known number of individuals is released, and $u_i(\mathbf{s}_j, t)$ is the likelihood an individual present at time $t$ and location $\mathbf{s}_j$. Conceptually, $\int_{\mathcal{A}} u_i(\mathbf{s}, t)d\mathbf{s}$ is the probability that an individual is present within the area $\mathcal{A}$, where $\mathcal{A} \subseteq \mathcal{S}$ and $\mathcal{S}$ is the study area. Similarly, $\theta_i \int_{\mathcal{A}} u_i(\mathbf{s}, t)d\mathbf{s}$ is the expected number of individuals of strain $i$ within the area $\mathcal{A}$. The $\frac{1}{|\mathcal{S}|}$ is a uniform probability density function over the study area $\mathcal{S}$ (i.e., $\int_{\mathcal{S}} \frac{1}{|\mathcal{S}|} d\mathbf{s} = 1$) and $\psi_i$ is a mixture probability.

The process $u_i(\mathbf{s}, t)$ is governed by the partial differential equation

$$\frac{\partial}{\partial t} u_i(\mathbf{s}, t) = \left( \frac{\partial^2}{\partial s_1^2} + \frac{\partial^2}{\partial s_2^2} \right) \mu_i(\mathbf{s}, t) u_i(\mathbf{s}, t) + \left( \frac{\partial}{\partial s_1} + \frac{\partial}{\partial s_2} \right) \nu_i(\mathbf{s}, t) u_i(\mathbf{s}, t) \tag{S3}$$

where $\mu_i(\mathbf{s}, t)$ and $\nu_i(\mathbf{s}, t)$ is the diffusion (or motility) rate and advection rate respectively. Both the diffusion and advection rate can vary spatially and/or temporally. In our study, we specify the diffusion rate using

$$\mu_i(\mathbf{s}, t) = \begin{cases} e^{\alpha_{i,0} + \alpha_{i,1} z(\mathbf{s}, t)} & \text{, if } x(t) = 1 \\ 0 & \text{, if } x(t) = 0 \end{cases}, \tag{S4}$$

where, for strain $i$, $\alpha_{i,0}$ is an intercept and $\alpha_{1,i}$ is a regression coefficient associated with temperature $z(\mathbf{s},t)$. The indicator variable $x(t)$ depends on the time, $t$, and is equal to 1 if it is daylight and equal to zero if it is night. Similarly we specified the advection rate as

$$\mu_i(\mathbf{s},t) = \begin{cases} \beta_{i,1} w(\mathbf{s},t) & \text{, if } x(t) = 1 \\ 0 & \text{, if } x(t) = 0 \end{cases}, \tag{S5}$$

where $\beta_{i,1}$ is a regression coefficient for wind velocity $w(\mathbf{s},t)$ for strain $i$.

Solving the PDE in Eq. S3 requires the specification of initial and boundary conditions. For initial conditions we used

$$u_i(\mathbf{s},t) = \begin{cases} 1 & \text{, if } \mathbf{s} = \boldsymbol{\omega}_i \\ 0 & \text{, if } \mathbf{s} \neq \boldsymbol{\omega}_i \end{cases}, \tag{S5}$$

where $\boldsymbol{\omega} = (0,0)'$ is the coordinate vector identifying the location the flies were released. We assumed Dirichlet boundary conditions and set $u(\mathbf{s},t) = 0$ at the edge of the study area.

Finally, to fully specify our Bayesian models we must select priors for all unknown parameters. For priors we used $\alpha_{i,0} \sim \text{unif}(-10,1)$, $\alpha_{i,1} \sim \text{unif}(-1,4)$, $\beta_{i,1} \sim \text{unif}(-0.05,0.05)$, $\psi_i \sim \text{unif}(0,1)$, $\phi_i \sim \text{unif}(0,100)$, and $p_i \sim \text{unif}(0,1)$. The priors for parameters associated with the PDE were chosen to result in a reasonable range of the diffusion rate (Eq. S4) and advection rate (Eq. S5) which was determined by viewing the resulting values of $u_i(\mathbf{s},t)$ obtained from simulation from the PDE in Eq. S3. The remaining parameters $\psi_i$, $\phi_i$, and $p_i$ were not associated with the PDE. For these parameters, we choose the priors to be "vague" or "non-informative." The parameters $\theta_i$ and $\boldsymbol{\omega}_i$, which correspond to the number of individuals released and the location of release where known in this study and therefore do not require priors.

#### 3.2 Required functions

Fitting the statistical model from section 3.2 to the experimental data is rather involved and required building an emulator to increase computational efficiency of the Marko chain Monte Carlo (MCMC) algorithm. Readers interested in understanding the details should consult Hooten and Hefley (2019; pgs. 501–515) and Hooten et al. (2011). For model implementation, we built seven functions given in the code below.

```r
# First-order neighborhood matrix from a RasterLayer object
neighborhood <- function(raster){
  nn <- matrix(,length(raster[]),4)
  for(i in 1:dim(nn)[1]){
        loc <- adjacent(raster,i)[,2]
        ln <- loc[which((loc+1)==i)]
        rn <- loc[which((loc-1)==i)]
        bn <- loc[which((loc-dim(raster)[2])==i)]
        tn <- loc[which((loc+dim(raster)[2])==i)]
        nn[i,1] <- if(length(ln)>0){ln}else{0}
        nn[i,2] <- if(length(rn)>0){rn}else{0}
        nn[i,3] <- if(length(bn)>0){bn}else{0}
        nn[i,4] <- if(length(tn)>0){tn}else{0}
  }
nn
}

# Propagator matrix to approximate PDE
propagator <- function(NN,x,mu,nu_ew,nu_ns,dx,dy,dt){
    T <- dim(mu)[2]
    Ht <- list()
  for(t in 1:T){
    if(x[t]==0){H <- diag(1,dim(NN)[1])}else{
    H <- matrix(0,dim(NN)[1],dim(NN)[1])
  for(i in 1:dim(H)[1]){
    if(length(which(NN[i,]>0))==4){
      H[i,i] <- 1-2*mu[i,t]*(dt/dx^2 + dt/dy^2)
      H[i,NN[i,1]] <- (dt/dx^2*mu[NN[i,1],t] - dt/dx*nu_ew[NN[i,1],t]/2)
      H[i,NN[i,2]] <- (dt/dx^2*mu[NN[i,2],t] + dt/dx*nu_ew[NN[i,2],t]/2)
      H[i,NN[i,3]] <- (dt/dy^2*mu[NN[i,3],t] - dt/dy*nu_ns[NN[i,3],t]/2)
      H[i,NN[i,4]] <- (dt/dy^2*mu[NN[i,4],t] + dt/dy*nu_ns[NN[i,4],t]/2)}
  }}
    Ht <- c(Ht,list(H))
  }
    Ht
```

```r
}

# RasterStack of u(s,t)
calc.u <- function(H,u0,t.stop,t.keep,ts){
  u.all <- u0
  u.all[] <- H[[1]]%*%u0[]
  u.all <- stack(mget(rep("u.all",t.stop)))
  for(t in 2:t.stop){
    H.t <- H[[t]]
    u.t <- u.all[[t-1]][]
    for(i in 1:ts){u.t <- H.t%*%u.t}
    u.all[[t]][] <- u.t
  }
  stack(u0,u.all)[[t.keep+1]]
}

#  Approximate PDE and return u(s,t)
u.sample <- function(y,x,z,w_ew,w_ns,omega,t.stop,t.keep,ts,dt,NN,alpha,beta,return){
  u0 <- raster(vals=0,ext=extent(z),res=res(z),crs=NA)
  u0[extract(u0,SpatialPoints(t(as.matrix(omega))),cellnumbers=TRUE)[1]] <- 1
  mu <- exp(alpha[1] + alpha[2]*z)
  nu_ew <- beta[1]*w_ew
  nu_ns <- beta[1]*w_ns
  H <- propagator(NN,x,mu[],nu_ew[],nu_ns[],res(mu)[1],res(mu)[2],dt)
  u.all <- calc.u(H,u0,t.stop,t.keep,ts)
  if(class(y)=="RasterStack"){vec(u.all[])[!is.na(vec(y[]))]}else{u.all}
}

# Get emulated u(s,t) for a given alpha and beta
ru <- function(UD,v.model,v.residuals,alpha,beta){
  parameters.pred <- data.frame(alpha0 = alpha[1],alpha1 = alpha[2],beta1 = beta[1])
  v.hat <- t(sapply(v.model,FUN=predict,newdata=parameters.pred,n.trees = v.model[[1]]$n.trees))
  e <- t(sapply(v.residuals,FUN=sample,size=1))
  v <- v.hat + e
  pnorm(UD%*%t(v))
}

# log likelihood using first order emulator
ll.em <- function(y,u,psi,theta,p,phi){
  p <- rep(p,each=227)
  y.obs <- vec(y[])[!is.na(vec(y[]))]
  sum(dnbinom(y.obs, size=1/phi, mu = theta*p*(psi*u+(1-psi)*(5^2/115^2)),log=TRUE))
}

# MCMC algorithm (see pgs. 505-511 in Hooten and Hefley (2019) for details)
mcmc.em <- function(y,UD,v.model,v.residuals,theta,
                    alpha.start,beta.start,psi.start,phi.start,p.start,
                    alpha.tune,beta.tune,psi.tune,phi.tune,p.tune,
                    n.mcmc){

  ####
  #### Setup Variables
  ####
  alpha.save <- matrix(,n.mcmc,2)
  beta.save <- matrix(,n.mcmc,1)
  psi.save <- matrix(,n.mcmc,1)
  phi.save <- matrix(,n.mcmc,1)
  p.save <- matrix(,n.mcmc,1)
  ll.save <- matrix(,n.mcmc,1)

  ####
  #### Starting values
  ####
  alpha <- alpha.start; alpha.cov <- matrix(c(1/10,-0.9/3,-0.9/3,1),2,2)
```

```
beta <- beta.start
psi <- psi.start
phi <- phi.start
p <- p.start

####
#### MCMC loop
####
for(k in 1:n.mcmc){

  ###
  ### Sample alpha
  ###
  alpha.star <- mvrnorm(1,alpha,alpha.tune*alpha.cov)
  if(alpha.star[1] > -10 & alpha.star[1] < 1 & alpha.star[2] > -1 & alpha.star[2] < 4){
  u <- ru(UD,v.model,v.residuals,alpha,beta)
  u.star <- ru(UD,v.model,v.residuals,alpha.star,beta)
  mh1 <- ll.em(y,u.star,psi,theta,p,phi)
  mh2 <- ll.em(y,u,psi,theta,p,phi)
  mh <- exp(mh1-mh2)}else{mh <- 0}
  if(mh > runif(1)){alpha <- alpha.star}

  ###
  ### Sample beta
  ###
  beta.star <- rnorm(1,beta,beta.tune)
  if(beta.star[1] > -0.05 & beta.star[1] < 0.05){
  u <- ru(UD,v.model,v.residuals,alpha,beta)
  u.star <- ru(UD,v.model,v.residuals,alpha,beta.star)
  mh1 <- ll.em(y,u.star,psi,theta,p,phi)
  mh2 <- ll.em(y,u,psi,theta,p,phi)
  mh <- exp(mh1-mh2)}else{mh <- 0}
  if(mh > runif(1)){beta <- beta.star}

  ###
  ### Sample psi
  ###
  psi.star <- rnorm(1,psi,psi.tune)
  if(psi.star > 0 & psi.star < 1){
    u <- ru(UD,v.model,v.residuals,alpha,beta)
    mh1 <- ll.em(y,u,psi.star,theta,p,phi)
    mh2 <- ll.em(y,u,psi,theta,p,phi)
    mh <- exp(mh1-mh2)}else{mh <- 0}
  if(mh > runif(1)){psi <- psi.star}

  ###
  ### Sample phi
  ###
  phi.star <- rnorm(1,phi,phi.tune)
  if(phi.star > 0 & phi.star < 100){
  u <- ru(UD,v.model,v.residuals,alpha,beta)
  mh1 <- ll.em(y,u,psi,theta,p,phi.star)
  mh2 <- ll.em(y,u,psi,theta,p,phi)
  mh <- exp(mh1-mh2)}else{mh <- 0}
  if(mh > runif(1)){phi <- phi.star}

  ###
  ### Sample p
  ###
  p.star <- rnorm(1,p,p.tune)
  if(p.star > 0 & p.star < 1){
    u <- ru(UD,v.model,v.residuals,alpha,beta)
    mh1 <- ll.em(y,u,psi,theta,p.star,phi)
    mh2 <- ll.em(y,u,psi,theta,p,phi)
```

```
      mh <- exp(mh1-mh2)}else{mh <- 0}
    if(mh > runif(1)){p <- p.star}

    ###
    ### Save samples & print interation number
    ###
    alpha.save[k,] <- alpha
    beta.save[k,] <- beta
    psi.save[k,] <- psi
    phi.save[k,] <- phi
    p.save[k,] <- p
    ll.save[k,] <- ll.em(y,u,psi,theta,p,phi)
    if(k%%100==0){print(k)}

    ###
    ### Adaptive random walk M-H algorithm (see pg. 238-239 in Computational Statistics by Given & Hoeting)
    ###
    T <- 100
    if(k%%T==0){
      alpha.cov <- cov(alpha.save[1:k,]) + diag(1/10^6,dim(alpha.cov)[1])
      accept.rate <- length(which(abs(diff(alpha.save[(T*(k/T-1)+1):(k),1]))>0))/T
      alpha.tune <- alpha.tune*exp(ifelse(accept.rate > 0.23,min(0.01,1/sqrt(k/T)),-min(0.01,1/sqrt(k/T))))^2
      accept.rate <- length(which(abs(diff(beta.save[(T*(k/T-1)+1):(k),1]))>0))/T
      beta.tune <- beta.tune*exp(ifelse(accept.rate > 0.44,min(0.01,1/sqrt(k/T)),-min(0.01,1/sqrt(k/T))))^2
      accept.rate <- length(which(abs(diff(psi.save[(T*(k/T-1)+1):(k),1]))>0))/T
      psi.tune <- psi.tune*exp(ifelse(accept.rate > 0.44,min(0.01,1/sqrt(k/T)),-min(0.01,1/sqrt(k/T))))^2
      accept.rate <- length(which(abs(diff(phi.save[(T*(k/T-1)+1):(k),1]))>0))/T
      phi.tune <- phi.tune*exp(ifelse(accept.rate > 0.44,min(0.01,1/sqrt(k/T)),-min(0.01,1/sqrt(k/T))))^2
      accept.rate <- length(which(abs(diff(p.save[(T*(k/T-1)+1):(k),1]))>0))/T
      p.tune <- p.tune*exp(ifelse(accept.rate > 0.44,min(0.01,1/sqrt(k/T)),-min(0.01,1/sqrt(k/T))))^2
      print("adapt proposals")
    }
  }

  ###
  ### Write output
  ###
  list(alpha=alpha.save,beta=beta.save,psi=psi.save,phi=phi.save,p=p.save,ll=ll.save)
}
```

##### 3.3 Emulator

Because the PDE component of our statistical model is computationally burdensome, we constructed an emulator using the approach outlined by Hooten et al. (2011). This can be implemented using the code below. This code takes approximately 10 hours to run using a desktop computer with optimized basic linear algebra subprograms, a 10-core 3.0 GHz processor and 128 GB of RAM. To reduce the time required to run the code, the number of times the PDE is solved can be reduced by changing `n.emulate` in the code below. Although this will make the code run faster, smaller values of `n.emulate`s may produce unreliable results.

```
# Preliminary steps
y <- rl.capture.rover
NN <- neighborhood(y[[1]])
t.keep <- unique(df.weather$Hours)[na.omit(match(unique(df.flies$Hours), unique(df.weather$Hours)))]/0.25
t.stop <- max(t.keep)
omega <- c(0, 0)  # Coordinates where flies where released
z <- ((rl.tempout - mean(rl.tempout[]))/sd(rl.tempout[]))
w_ew <- rl.wind_ew
w_ns <- rl.wind_ns
x <- df.weather$Daylight

# Step 1 & 2 on pg 483 of Hooten et al. (2011)
n.emulate <- 10000
u.emulate <- matrix(, sum(!is.na(vec(y[]))), n.emulate)
alpha.emulate <- matrix(, 2, n.emulate)
```

```
beta.emulate <- matrix(, 1, n.emulate)

set.seed(4222)
ptm <- proc.time()
for (i in 1:n.emulate) {
    a <- pnorm(mvrnorm(n = 1, rep(0, 2), Sigma = matrix(c(1, -0.7, -0.7, 1), 2, 2)))
    alpha.emulate[, i] <- c(a[1] * 11 - 10, a[2] * 5 - 1)
    beta.emulate[, i] <- runif(1, -0.05, 0.05)
    u.emulate[, i] <- u.sample(y = y, x = x, z = z, w_ew = w_ew, w_ns = w_ns, omega = omega,
        t.stop = t.stop, t.keep = t.keep, ts = 45, dt = 1/3, NN = NN, alpha = alpha.emulate[,
            i], beta = beta.emulate[, i])
    print(i)
}
proc.time() - ptm

# Step 3 on pg 483 of Hooten et al. (2011)
keep <- which(apply(u.emulate, 2, min) >= 0 & apply(u.emulate, 2, max) < 1)
Y <- qnorm(u.emulate[, keep])
D <- diag(svd(Y)$d)
U <- svd(Y)$u
V <- svd(Y)$v
UD <- (U %*% D)[, 1:5]  # dimension reduction
V <- V[, 1:5]  # dimension reduction

# Step 4 on pg 483 of Hooten et al. (2011)
v.model <- list()
v.residuals <- list()
parameters <- data.frame(alpha0 = alpha.emulate[1, keep], alpha1 = alpha.emulate[2, keep],
    beta1 = beta.emulate[1, keep])
for (i in 1:dim(V)[2]) {
    df.temp <- data.frame(v = V[, i], parameters)
    m <- gbm(v ~ alpha0 + alpha1 + beta1, data = df.temp, distribution = "laplace", interaction.depth = 3,
        n.trees = 2000, shrinkage = 0.1, bag.fraction = 1)
    e <- df.temp$v - predict(m, n.trees = m$n.trees)
    v.model[i] <- list(m)
    v.residuals[i] <- list(e)
    print(i)
}
```

#### 3.4 Model fitting (sitters)

Below is the code that uses an MCMC algorithm to fit the Bayesian model from section 3.1 to the experimental data for sitters. We draw 50,000 samples in the code below. To reduce the time required to run the code, the number of samples drawn can be reduced by specifying `n.mcmc` to be a smaller number, however, this may may produce unreliable results. As with all MCMC algorithms, it is important to check to ensure that the Markov chains are efficiently sampling from the stationary distribution. Below is code that can be used to examine the trace plots for all parameters.

```
set.seed(1422)
ptm <- proc.time()
samples.sitter <- mcmc.em(y = rl.capture.sitter, UD = UD, v.model = v.model, v.residuals = v.residuals,
    theta = 5352, alpha.start = c(-2, 0), beta.start = c(0), psi.start = 0.5, phi.start = 2.5,
    p.start = 0.1, alpha.tune = 1/5, beta.tune = 3/1000, psi.tune = 5/100, phi.tune = 5/10,
    p.tune = 5/100, n.mcmc = 50000)
proc.time() - ptm

# Check trace plots
plot(samples.sitter$alpha[, 1], typ = "l")
plot(samples.sitter$alpha[, 2], typ = "l")
plot(samples.sitter$beta[, 1], typ = "l")
plot(samples.sitter$psi[, 1], typ = "l")
plot(samples.sitter$phi[, 1], typ = "l")
plot(samples.sitter$p[, 1], typ = "l")
plot(samples.sitter$ll[, 1], typ = "l")
```

#### 3.5 Model fitting (rovers)

Below is the code that uses an MCMC algorithm to fit the Bayesian model from section 3.1 to the experimental data for rovers. We draw 50,000 samples in the code below. To reduce the time required to run the code, the number of samples drawn can be reduced by specifying `n.mcmc` to be a smaller number, however, this may may produce unreliable results. As with all MCMC algorithms, it is important to check to ensure that the Markov chains are efficiently sampling from the stationary distribution. Below is code that can be used to examine the trace plots for all parameters.

```
set.seed(5322)
ptm <- proc.time()
samples.rover <- mcmc.em(y = rl.capture.rover, UD = UD, v.model = v.model, v.residuals = v.residuals,
    theta = 5644, alpha.start = c(-3, 0), beta.start = c(0), psi.start = 0.5, phi.start = 2.5,
    p.start = 0.1, alpha.tune = 1/5, beta.tune = 3/1000, psi.tune = 5/100, phi.tune = 5/10,
    p.tune = 5/100, n.mcmc = 50000)
proc.time() - ptm

# Check trace plots
burn.in <- 1
plot(samples.rover$alpha[, 1], typ = "l")
plot(samples.rover$alpha[, 2], typ = "l")
plot(samples.rover$beta[, 1], typ = "l")
plot(samples.rover$psi[, 1], typ = "l")
plot(samples.rover$phi[, 1], typ = "l")
plot(samples.rover$p[, 1], typ = "l")
plot(samples.rover$ll[, 1], typ = "l")
```

#### 3.6 Model fitting (outbred)

Below is the code that uses an MCMC algorithm to fit the Bayesian model from section 3.1 to the experimental data for the outbred strain. We draw 50,000 samples in the code below. To reduce the time required to run the code, the number of samples drawn can be reduced by specifying `n.mcmc` to be a smaller number, however, this may may produce unreliable results. As with all MCMC algorithms, it is important to check to ensure that the Markov chains are efficiently sampling from the stationary distribution. Below is code that can be used to examine the trace plots for all parameters.

```
set.seed(5292)
ptm <- proc.time()
samples.sudbury <- mcmc.em(y = rl.capture.sudbury, UD = UD, v.model = v.model, v.residuals = v.residuals,
    theta = 5657, alpha.start = c(-3, 0), beta.start = c(0), psi.start = 0.5, phi.start = 2.5,
    p.start = 0.1, alpha.tune = 1/5, beta.tune = 3/1000, psi.tune = 5/100, phi.tune = 5/10,
    p.tune = 5/100, n.mcmc = 50000)
proc.time() - ptm

# Check trace plots
burn.in <- 1
plot(samples.sudbury$alpha[, 1], typ = "l")
plot(samples.sudbury$alpha[, 2], typ = "l")
plot(samples.sudbury$beta[, 1], typ = "l")
plot(samples.sudbury$psi[, 1], typ = "l")
plot(samples.sudbury$phi[, 1], typ = "l")
plot(samples.sudbury$p[, 1], typ = "l")
plot(samples.sudbury$ll[, 1], typ = "l")
```

#### 3.7 Derived quantities

In the manuscript several derived quantities (i.e., functions of the posterior distribution) were mentioned. Below is the R code to obtain the posterior mean of the derived quantities.

```
# Difference in diffusion over 24 hours for rovers vs. outbred at 22 celsius
temp <- 22  # Temperature in celsius
z <- (temp - mean(rl.tempout[]))/sd(rl.tempout[])
dq <- 1440 * (exp(samples.rover$alpha[, 1] + samples.rover$alpha[, 2] * z) - exp(samples.sudbury$alpha[,
    1] + samples.sudbury$alpha[, 2] * z))
mean(dq)
```

```
# Difference in diffusion over 24 hours for sitters vs. outbred at 22 celsius
temp <- 22  # Temperature in celsius
z <- (temp - mean(rl.tempout[]))/sd(rl.tempout[])
dq <- 1440 * (exp(samples.sitter$alpha[, 1] + samples.sitter$alpha[, 2] * z) - exp(samples.sudbury$alpha[,
    1] + samples.sudbury$alpha[, 2] * z))
mean(dq)

# Difference in diffusion over 24 hours for rovers vs. outbred at 14 celsius
temp <- 14  # Temperature in celsius
z <- (temp - mean(rl.tempout[]))/sd(rl.tempout[])
dq <- 1440 * (exp(samples.rover$alpha[, 1] + samples.rover$alpha[, 2] * z) - exp(samples.sudbury$alpha[,
    1] + samples.sudbury$alpha[, 2] * z))
mean(dq)

# Difference in diffusion over 24 hours for sitters vs. outbred at 14 celsius
temp <- 14  # Temperature in celsius
z <- (temp - mean(rl.tempout[]))/sd(rl.tempout[])
dq <- 1440 * (exp(samples.sitter$alpha[, 1] + samples.sitter$alpha[, 2] * z) - exp(samples.sudbury$alpha[,
    1] + samples.sudbury$alpha[, 2] * z))
mean(dq)

# Difference in advection over 24 hours for sitters vs. rovers at 1 meter per second wind velocity
wind <- 1  # wind velocity  meters per second
dq <- 1440 * (samples.sitter$beta[, 1] * wind - samples.rover$beta[, 1] * wind)
mean(dq)

# Difference in advection over 24 hours for sitters vs. rovers at 2.5 meter per second wind velocity
wind <- 2.5  # wind velocity  meters per second
dq <- 1440 * (samples.sitter$beta[, 1] * wind - samples.rover$beta[, 1] * wind)
mean(dq)
```

##### 3.8 Figure 4

Below is R code that can be used to reproduce Figure 4.

```
burn.in <- 10000
samples.alpha0 <- cbind(samples.sitter$alpha[-c(1:burn.in), 1], samples.rover$alpha[-c(1:burn.in),
    1], samples.sudbury$alpha[-c(1:burn.in), 1])
alpha0.ev <- apply(samples.alpha0, 2, mean)
alpha0.ui <- apply(samples.alpha0, 2, quantile, prob = 0.975)
alpha0.li <- apply(samples.alpha0, 2, quantile, prob = 0.025)
samples.alpha1 <- cbind(samples.sitter$alpha[-c(1:burn.in), 2], samples.rover$alpha[-c(1:burn.in),
    2], samples.sudbury$alpha[-c(1:burn.in), 2])
alpha1.ev <- apply(samples.alpha1, 2, mean)
alpha1.ui <- apply(samples.alpha1, 2, quantile, prob = 0.975)
alpha1.li <- apply(samples.alpha1, 2, quantile, prob = 0.025)
samples.beta1 <- cbind(samples.sitter$beta[-c(1:burn.in), 1], samples.rover$beta[-c(1:burn.in),
    1], samples.sudbury$beta[-c(1:burn.in), 1])
beta1.ev <- apply(samples.beta1, 2, mean)
beta1.ui <- apply(samples.beta1, 2, quantile, prob = 0.975)
beta1.li <- apply(samples.beta1, 2, quantile, prob = 0.025)
samples.psi <- cbind(samples.sitter$psi[-c(1:burn.in), 1], samples.rover$psi[-c(1:burn.in),
    1], samples.sudbury$psi[-c(1:burn.in), 1])
psi.ev <- apply(samples.psi, 2, mean)
psi.ui <- apply(samples.psi, 2, quantile, prob = 0.975)
psi.li <- apply(samples.psi, 2, quantile, prob = 0.025)

# pdf(file='Fig_4.pdf',width=6,height=2,onefile=FALSE)
par(mfrow = c(1, 3))
par(mar = c(6, 4, 2, 2))
plotCI(1:3, alpha0.ev, ui = alpha0.ui, li = alpha0.li, pch = 20, xlab = "", xaxt = "n", ylab = expression(alpha[0]),
    las = 2, main = "(a)")
axis(1, a = 1:3, labels = c("Sitter", "Rover", "Outbred"), las = 2)
plotCI(1:3, alpha1.ev, ui = alpha1.ui, li = alpha1.li, pch = 20, xlab = "", xaxt = "n", ylab = expression(alpha[1]),
```

```
    las = 2, main = "(b)")
axis(1, a = 1:3, labels = c("Sitter", "Rover", "Outbred"), las = 2)
plotCI(1:3, beta1.ev, ui = beta1.ui, li = beta1.li, pch = 20, xlab = "", xaxt = "n", ylab = expression(beta[1]),
    las = 2, main = "(c)")
axis(1, a = 1:3, labels = c("Sitter", "Rover", "Outbred"), las = 2)
# dev.off()
```

#### 3.9 Figure 5

Below is R code that can be used to reproduce Figure 5.

```
burn.in <- 10000
n.samples <- dim(samples.sitter$alpha)[1]
temp <- seq(10, 22.5, by = 0.1)
z <- (temp - mean(rl.tempout[]))/sd(rl.tempout[])
diffusion.sitter <- matrix(, length(z), n.samples - burn.in)
diffusion.rover <- matrix(, length(z), n.samples - burn.in)
diffusion.sudbury <- matrix(, length(z), n.samples - burn.in)

for (i in (burn.in + 1):n.samples) {
    alpha <- samples.sitter$alpha[i, 1:2]
    diffusion.sitter[, i - burn.in] <- exp(alpha[1] + alpha[2] * z)

    alpha <- samples.rover$alpha[i, 1:2]
    diffusion.rover[, i - burn.in] <- exp(alpha[1] + alpha[2] * z)

    alpha <- samples.sudbury$alpha[i, 1:2]
    diffusion.sudbury[, i - burn.in] <- exp(alpha[1] + alpha[2] * z)
}

# pdf(file='Fig_5.pdf',width=6,height=6,onefile=FALSE)
par(mfrow = c(1, 1))
par(mar = c(4, 5, 2, 2))
plot(temp, rowMeans(diffusion.sitter), typ = "l", lwd = 3, ylim = c(0, 0.3), xlab = expression("Temperature(" *
    degree * C * ")"), ylab = expression("Diffusion rate(" * m^2 * min^-1 * ")"))
lwr.CI <- apply(diffusion.sitter, 1, FUN = quantile, prob = c(0.025))
upper.CI <- apply(diffusion.sitter, 1, FUN = quantile, prob = c(0.975))
polygon(c(temp, rev(temp)), c(lwr.CI, rev(upper.CI)), col = adjustcolor("black", alpha.f = 0.1),
    border = NA)

points(temp, rowMeans(diffusion.rover), typ = "l", lwd = 3, col = "deepskyblue")
lwr.CI <- apply(diffusion.rover, 1, FUN = quantile, prob = c(0.025))
upper.CI <- apply(diffusion.rover, 1, FUN = quantile, prob = c(0.975))
polygon(c(temp, rev(temp)), c(lwr.CI, rev(upper.CI)), col = adjustcolor("deepskyblue", alpha.f = 0.1),
    border = NA)

points(temp, rowMeans(diffusion.sudbury), typ = "l", lwd = 3, col = "indianred")
lwr.CI <- apply(diffusion.sudbury, 1, FUN = quantile, prob = c(0.025))
upper.CI <- apply(diffusion.sudbury, 1, FUN = quantile, prob = c(0.975))
polygon(c(temp, rev(temp)), c(lwr.CI, rev(upper.CI)), col = adjustcolor("indianred", alpha.f = 0.1),
    border = NA)
rug(df.weather$tempOut)
legend(x = 10, y = 0.3, cex = 1.3, legend = c("Sitter", "Rover", "Outbred"), bty = "n", lty = 1,
    lwd = 2, col = c("black", "deepskyblue", "indianred"))
# dev.off()
```

#### 3.10 Figure 6

Below is R code that can be used to reproduce Figure 6.

```
burn.in <- 10000
n.samples <- dim(samples.sitter$beta)[1]
w <- seq(0, 2.7, by = 0.01)
```

```r
advection.sitter <- matrix(, length(w), n.samples - burn.in)
advection.rover <- matrix(, length(w), n.samples - burn.in)
advection.sudbury <- matrix(, length(w), n.samples - burn.in)

for (i in (burn.in + 1):n.samples) {
    beta <- samples.sitter$beta[i, 1]
    advection.sitter[, i - burn.in] <- beta[1] * w

    beta <- samples.rover$beta[i, 1]
    advection.rover[, i - burn.in] <- beta[1] * w

    beta <- samples.sudbury$beta[i, 1]
    advection.sudbury[, i - burn.in] <- beta[1] * w
}

# pdf(file='Fig_6.pdf',width=6,height=6,onefile=FALSE)
par(mfrow = c(1, 1))
par(mar = c(4, 5, 2, 2))
plot(w, rowMeans(advection.sitter), typ = "l", lwd = 3, ylim = c(0,
    0.1), xlab = expression("Wind speed (meters per second)"),
    ylab = expression("Advection rate (meters per min)"))
lwr.CI <- apply(advection.sitter, 1, FUN = quantile, prob = c(0.025))
upper.CI <- apply(advection.sitter, 1, FUN = quantile, prob = c(0.975))
polygon(c(w, rev(w)), c(lwr.CI, rev(upper.CI)), col = adjustcolor("black",
    alpha.f = 0.1), border = NA)

points(w, rowMeans(advection.rover), typ = "l", lwd = 3, col = "deepskyblue")
lwr.CI <- apply(advection.rover, 1, FUN = quantile, prob = c(0.025))
upper.CI <- apply(advection.rover, 1, FUN = quantile, prob = c(0.975))
polygon(c(w, rev(w)), c(lwr.CI, rev(upper.CI)), col = adjustcolor("deepskyblue",
    alpha.f = 0.1), border = NA)

points(w, rowMeans(advection.sudbury), typ = "l", lwd = 3, col = "indianred")
lwr.CI <- apply(advection.sudbury, 1, FUN = quantile, prob = c(0.025))
upper.CI <- apply(advection.sudbury, 1, FUN = quantile, prob = c(0.975))
polygon(c(w, rev(w)), c(lwr.CI, rev(upper.CI)), col = adjustcolor("indianred",
    alpha.f = 0.1), border = NA)
rug(df.weather$WindSpeed)
legend(x = 0, y = 0.1, cex = 1.3, legend = c("Sitter", "Rover",
    "Outbred"), bty = "n", lty = 1, lwd = 2, col = c("black",
    "deepskyblue", "indianred"))
# dev.off()
```

### References

Edelsparre, A. H., Hefley, T. J., Rodriguez, M. A., Fitzpatrick, M. J., and Sokolowski, M. B. (2020). Data for: Scaling up: understanding movement from individual differences to population-level dispersal. .

Givens, G. H. and Hoeting, J. A. (2012). *Computational Statistics*. John Wiley & Sons.

Hooten, M. B. and Hefley, T. J. (2019). *Bringing Bayesian Models to Life*. Chapman & Hall/CRC.

Hooten, M. B., Leeds, W. B., Fiechter, J., and Wikle, C. K. (2011). Assessing first-order emulator inference for physical parameters in nonlinear mechanistic models. *Journal of Agricultural, Biological, and Environmental Statistics*, 16(4):475–494.

Wikle, C. K., Zammit-Mangion, A., and Cressie, N. (2019). *Spatio-temporal Statistics with R*. CRC Press.
